# Injectable Liposome-based Supramolecular Hydrogels for the Programmable Release of Multiple Protein Drugs

**DOI:** 10.1101/2021.09.26.461871

**Authors:** Santiago Correa, Abigail K. Grosskopf, John H. Klich, Hector Lopez Hernandez, Eric A. Appel

**Affiliations:** Department of Materials Science & Engineering, Stanford University, Stanford, CA 94305, USA; Department of Chemical Engineering, Stanford University, Stanford, CA 94305, USA; Department of Bioengineering, Stanford University, Stanford, CA 94305, USA; ChEM-H Institute, Stanford University, Stanford, CA 94305, USA; Department of Pediatrics – Endocrinology, Stanford University School of Medicine, Stanford, CA 94305, USA; Woods Institute for the Environment, Stanford University, Stanford, CA 94305, USA

**Keywords:** Hydrogels, Liposomes, Rheology, Viscoelasticity, Controlled delivery, Proteins

## Abstract

Directing and manipulating biological functions is at the heart of next-generation biomedical initiatives such as tissue and immuno-engineering. Yet, the ambitious goal of engineering complex biological networks requires the ability to precisely perturb specific signaling pathways at distinct times and places. Using lipid nanotechnology and the principles of supramolecular self-assembly, we have developed an injectable liposomal nanocomposite hydrogel platform to precisely control drug presentation through programing of the co-release of multiple protein drugs. These liposomal hydrogels exhibited robust shear-thinning and self-healing behaviors enabling facile injectability for local drug delivery applications. By integrating modular lipid nanotechnology into this hydrogel platform, we introduced multiple mechanisms of protein release based on liposome surface chemistry. When injected into immuno-competent mice, these liposomal hydrogels exhibited formulation-dependent rates of dissolution and excellent biocompatibility. To fully validate the utility of this system for multi-protein delivery, we demonstrated the synchronized, sustained, and localized release of IgG antibody and IL-12 cytokine *in vivo*, despite the significant size differences between these two proteins. Overall, these liposomal nanocomposite hydrogels are a highly modular platform technology with the ability the mediate orthogonal modes of protein release and the potential to precisely coordinate biological cues both *in vitro* and *in vivo*.

## Introduction

Protein drugs are essential tools for engineering biological systems to improve human health. Challenging biomedical applications such as tissue regeneration and wound healing often rely on diverse growth factors and cytokines.^1,2^ Likewise, immunotherapy relies heavily on antibodies, engineered proteins, cytokines, and chemokines to activate or dampen the immune system (e.g., for cancer or autoimmune disorders, respectively).^3^ While tissue regeneration, wound healing, and immuno-engineering pursue different biological outcomes, they share a similar challenge – to manipulate complex biological networks. The capacity to successfully rewire these networks rests on the ability to perturb multiple dynamic signaling pathways at specific times^4^ and within specific tissues (e.g., diseased tissues or specific lymphoid organs) to achieve desired therapeutic outcomes.^5^ Indeed, disruption of these networks outside of the appropriate timeframe or outside of the target tissues can lead to serious side effects.^5,6^ Taken together, technology that can precisely coordinate the release kinetics of diverse protein drugs in specific locations are critical for the controlled modulation of biological systems.

Nanomedicine and injectable hydrogel technologies seek to provide precise spatiotemporal control over drug delivery.^6-8^ In terms of spatial control, nanoparticle drug carriers seek to preferentially accumulate in target tissues after systemic administration, but besides filtration organs like the liver and spleen, their ability to target to specific tissues is limited.^9-11^ In terms of scheduled multi-drug release, nanomedicine has been successful in staging the release of different classes of drugs (e.g., hydrophilic vs. hydrophobic small molecules, nucleic acids, proteins), but staged delivery of proteins remains a challenge.^8,12^ Meanwhile, the emergence of injectable hydrogels has provided a significant improvement for minimally invasive localized therapy,^13-22^ but scheduled multi-drug release from these systems is often limited to strategies that leverage size-governed (e.g., smaller drugs first) or solubility-governed (e.g., most soluble drugs first) release mechanisms.^7,23^ Overall, there remains no minimally invasive delivery technology that can tune the relative release rates of multiple protein drugs *in vivo*.

To address this technology gap, we developed a new supramolecular hydrogel platform constructed directly from dodecyl-modified hydroxypropylmethylcellulose (HPMC-C_12_) and liposomal nanoparticle building blocks (Scheme 1A). This hydrogel material was inspired by previously developed polymer-nanoparticle hydrogels based on other solid nanoparticles.^24-26^ By integrating liposomes directly into the hydrogel network, this system possesses several distinct drug compartments useful for multi-protein delivery. Moreover, as an injectable hydrogel, this system allows for highly localized delivery of cargo which is not possible with free liposomal carriers.

In this report, we provide a comprehensive assessment of how the clinically-relevant rheological properties of these liposomal nanocomposite hydrogels (LNHs) are influenced by critical formulation parameters (e.g., liposome size and concentration). In general, the mechanical properties of the LNHs are readily tuned by liposome concentration and to a lesser extent size, and they are robust to changes in liposome composition. We performed preclinical toxicology studies in immuno-competent mice to determine the biocompatibility of LNHs, where we observed a steady erosion of the hydrogel over time with no signs of toxicity. Finally, we determined the ability of LNHs to stage the release of protein drugs through orthogonal passive, electrostatic, and affinity-based release mechanisms (Scheme 2B). We validated the utility of our orthogonal release approach by using LNHs to synchronize the sustained local delivery of IgG antibody and IL-12 cytokines *in vivo*, despite the significant size difference between these two protein drugs. Overall, LNHs leverage the impressive capabilities of lipid nanotechnology to generate a shear-thinning and self-healing biomaterial platform capable of simultaneously accommodating numerous mechanisms of protein drug release. We anticipate this platform will provide a readily generalizable approach for the localized staged release of protein drugs with implications for the fields of immunotherapy and regenerative medicine.

**Scheme 1.**
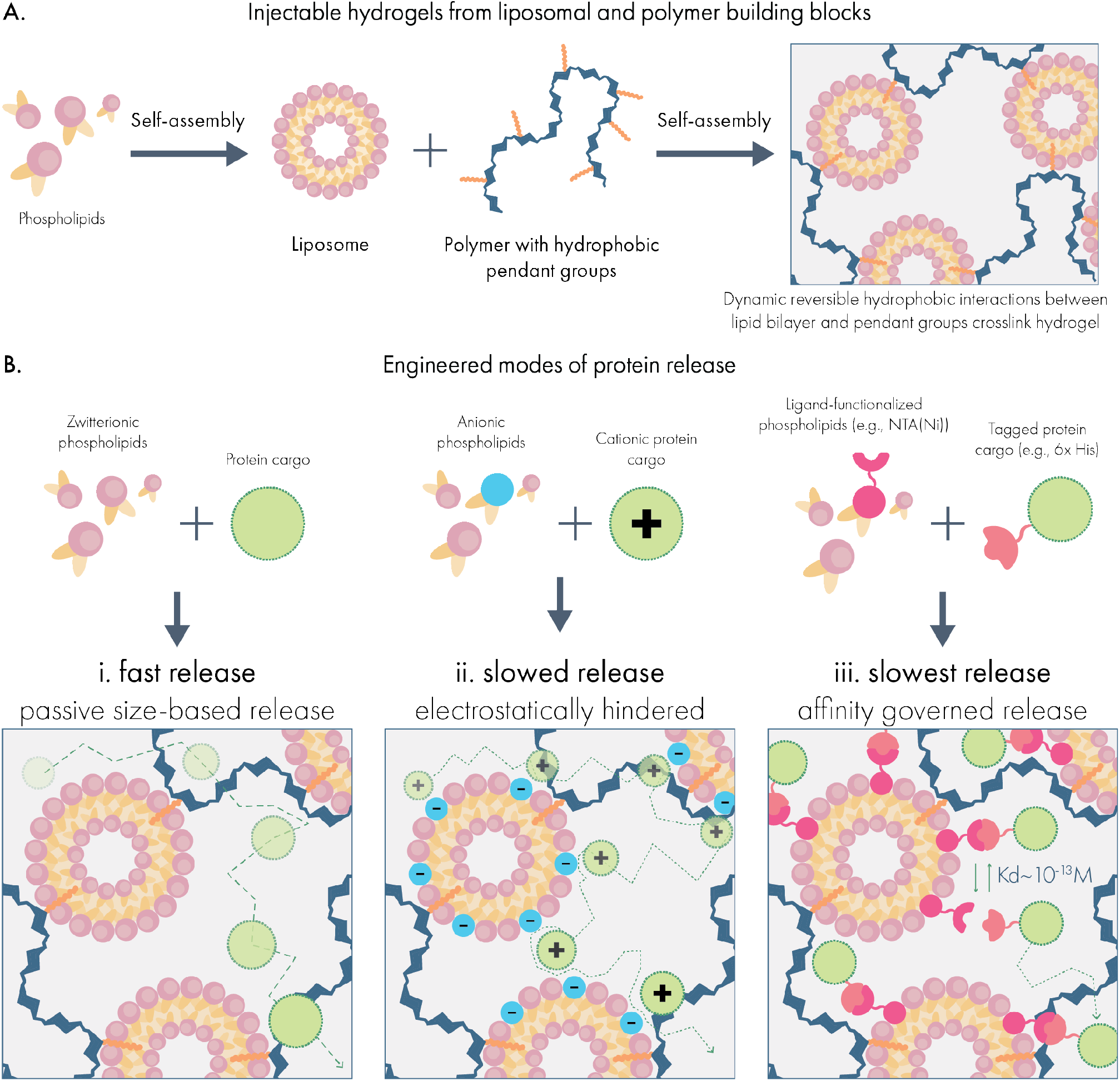
Forming injectable hydrogels for multi-modal protein drug delivery from liposomal building blocks. **(A)** Liposomes self-assemble from phospholipids to form nanoparticles featuring a hydrophobic compartment within the lipid bilayer. When liposomes are mixed with dodecyl-modified hydroxypropyl methylcellulose (HPMC-C_12_), the hydrophobic dodecyl pendants insert into the lipid membrane, generating a crosslinked hydrogel network. **(B)** Drug delivery capabilities are tuned by incorporating functional phospholipids into the liposomal building blocks, which establish interactions with protein cargo in the hydrogel. These include **(i)** zwitterionic liposomes that minimize interactions with cargo and mediate release by size-based diffusion; **(ii)** charge-carrying liposomes that engage with proteins exhibiting an opposite net charge; and **(iii)** ligand-functionalized liposomes that establish affinity-mediated release of specific protein cargo.

## Results and Discussion

### Liposomes spontaneously form injectable hydrogels when mixed with hydrophobically-modified biopolymers

To stably incorporate liposomal nanoparticles into a supramolecular hydrogel network, we exploited the ability for hydrophobic carbon chains to post-insert into lipid membranes.^27^ To this end, we modified high molecular weight hydroxypropyl methylcellulose with fatty dodecyl side chains (HPMC-C_12_) to introduce a molecular motif capable of post-insertion. Initially, we prepared liposomal nanoparticles composed of 1,2-dimyristoyl-sn-glycero-3-phosphocholine (DMPC; zwitterionic headgroup), 1,2-dimyristoyl-sn-glycero-3-phospho-(1’-rac-glycerol) (DMPG; anionic headgroup), and cholesterol at a 9:1:2 molar ratio using the thin-film rehydration and extrusion method. ^28^ This formulation was chosen for its previously reported utility for drug delivery.^28^ Following extrusion through 50 nm polycarbonate filters, liposomes exhibited a monodisperse average hydrodynamic size of 90.4 nm and a PDI of 0.1, indicating uniform nanoparticle preparation. Physical mixing of HPMC-C_12_ and liposome solutions led to the generation of a pearlescent supramolecular hydrogel comprising 2wt% polymer and 10wt% liposome, which we denote as 2P-10L.

Dynamic oscillatory shear rheology of the resulting 2P-10L hydrogel revealed robust solid-like characteristics (e.g., G’ greater than G’’) over the entire range of frequencies evaluated (Figure 1A). The robust mechanical properties of these LNHs, with no observable crossover of G’ and G’’ in the low frequency oscillation regime, indicates successful hydrogel formation and sets these materials apart from previously reported liposome-based hydrogels that exhibit significant viscoelasticity.^29^ In contrast, when liposomes were mixed with unmodified HPMC, no hydrogel was formed and rheological characterization revealed the mixture exhibited liquid-like properties (e.g., G’’ greater than G’) over physiologically relevant frequencies (Supplemental Figure 1B). Likewise, solutions of 2wt% HPMC-C_12_ fail to form robust hydrogels in the absence of liposomes (Supplemental Figure 1C). Overall, these data strongly indicate that LNHs are formed from multivalent and dynamic polymer-particle crosslinking interactions between the liposomal and polymer building blocks that requires the presence of a hydrophobic molecular motif on the cellulosic polymer component.

**Figure 1.**
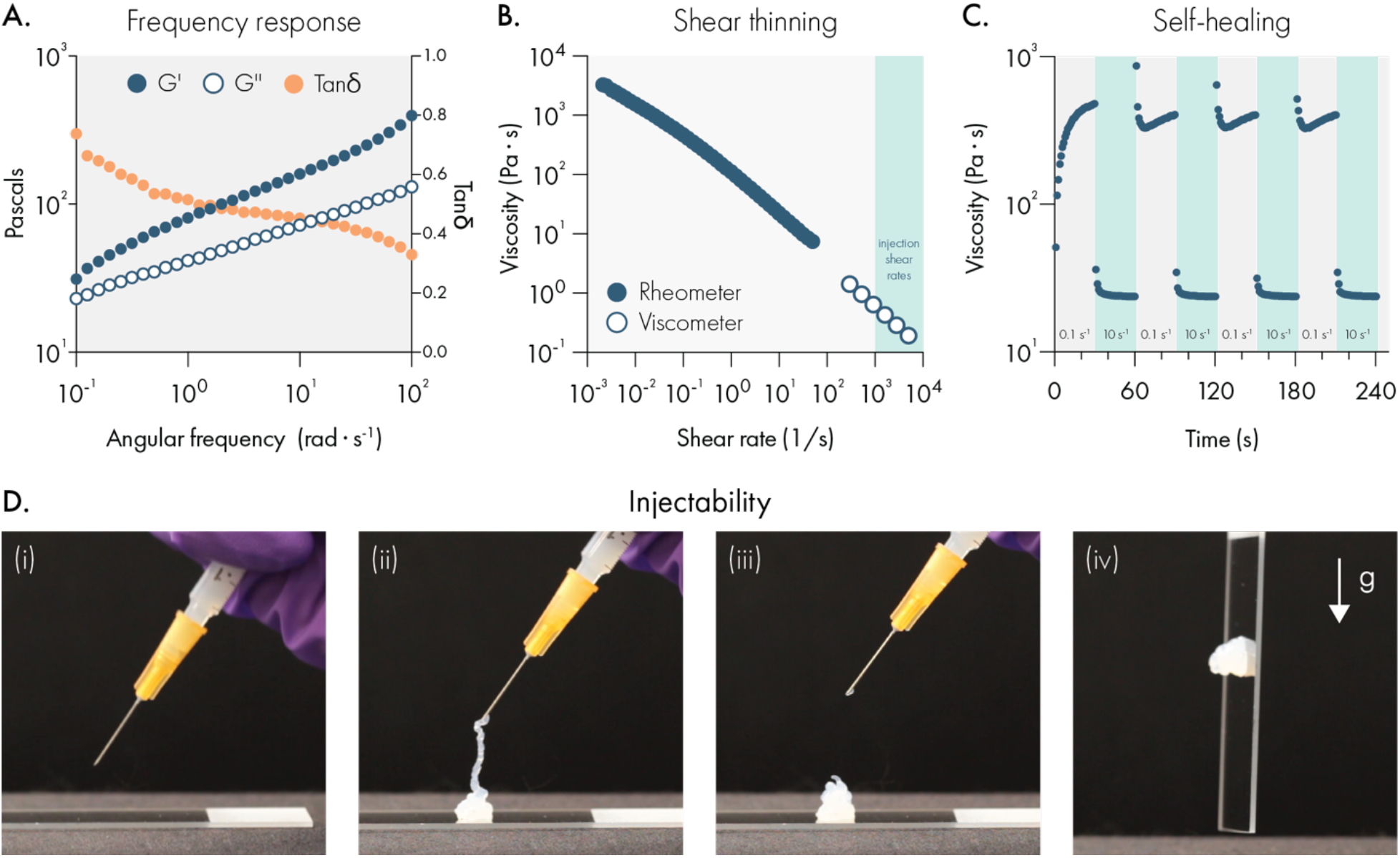
Liposomal hydrogels exhibit shear-thinning and self-healing mechanical properties that enable injection through needles. **(A)** Dynamic oscillatory shear rheology of liposomal hydrogels (2wt% HPMC-C_12_; 10wt% 50 nm extruded liposome) indicates the storage modulus (G’) is greater than the loss modulus (G’’) over a broad range of frequencies, demonstrating robust solid-like properties. **(B)** Steady shear rheology measurements (filled circles) and viscometer measurements (empty circles) indicate that the viscosity of liposomal hydrogels steadily decreases by roughly 5-orders of magnitude as shear rates approach injection conditions. **(C)** Step-shear rheology measurements demonstrate that liposomal hydrogels can repeatedly regain their original viscosity after exposure to high shear conditions. **(D)** Representative photographs of liposomal hydrogels injected through a 26-gauge needle fitted onto a 1 mL syringe.

We then set out to determine whether LNHs exhibited necessary mechanical properties for the development of a clinically-relevant drug delivery platform.^30^ First, we evaluated whether these materials possess shear-thinning and self-healing properties, which would allow them to serve as injectable depots of therapeutic compounds. To test this, we performed steady shear rheology and viscometry to evaluate how LNH viscosity changes under high shear conditions, representative of the shear forces in a syringe needle (Figure 1B). We observed that LNH viscosity drops approximately 4 orders of magnitude as shear rates increase to levels that simulate syringe-needle injection, demonstrating that this material is extremely injectable. The steep decrease in the viscosity suggests that the internal structure of the LNH network is disrupted during high-shear conditions, which would require the structure to spontaneously regenerate in order for the LNH to resolidify after injection.^30^ To determine the ability for LNHs to self-heal, we conducted step-shear experiments where the material undergoes multiple cycles of high and low shear conditions (Figure 1C). We observed the viscosity of LNHs rapidly and repeatedly restored to original levels once high-shear conditions were suspended. We also observed that LNHs could be easily injected through 1 mL syringes fitted with 26-gauge needles and re-solidify after injection (Figure 1D, Supplemental Video 1). These data indicate that LNHs are able to dynamically re-assemble the hydrogel network following its disruption, consistent with a supramolecular mechanism of assembly and providing a minimally invasive administration route for drug delivery.

Our next objective was to determine how robust these supramolecular assemblies were to changes in the liposome formulation. Our initial liposomes followed an anionic formulation previously validated for drug delivery,^28^ but one of the assets of lipid nanotechnology is the broad parameter space available to tune key properties (e.g., surface chemistry, surface charge, anti-fouling, and melting temperature) through careful selection of the individual phospholipid components. We evaluated how a cationic version of our liposomes (replacing the 1,2-dimyristoyl-sn-glycero-3-phosphoglycerol with the cationic phospholipid 1,2-dioleoyl-3-trimethylammonium-propane) behaved when mixed with HPMC-C_12_ and observed that these systems yielded hydrogels with similar rheological properties to our original formulations (Supplemental Figure 2). Like the hydrogels formed with anionic liposomes, the cationic liposomal hydrogels were solid-like over the same range of frequencies and were both shear-thinning and self-healing. Next, we evaluated how a PEGylated liposomal formulation behaved in this platform, and we observed overall consistent mechanical properties (Supplemental Figure 3). This compatibility with diverse liposomal formulations expands the biomedical capabilities of these materials. For example, the ability to incorporate cationic liposomes is promising for the development of injectable materials for gene delivery.^31^ Likewise, the anti-fouling capabilities of PEGylated formulations could be leveraged to reduce engagement with the host after injection.^32^ Moreover, since most clinically approved liposomal drugs feature PEGylation, this technology could provide a means to locally administer approved liposomal drugs through minimally invasive injections.

Having confirmed that anionic, cationic, and PEGylated liposomes yield consistent hydrogels in our system, we began to evaluate the parameters of cholesterol content and lipid melting temperature (T_m_) on the resulting rheological properties of these materials. These properties are often varied to tune drug retention and liposome stability.^33,34^ Using our anionic liposome formulation as a starting point, we evaluated hydrogels generated from liposomes composed of 17, 37.5, and 50 mol% cholesterol (Supplemental Figure 4). Overall, we observed that the modulus of the materials and the shear-thinning behaviors were consistent as cholesterol content was varied. Yet, we did observe a difference in the yield strain between high and low cholesterol formulations, with low-cholesterol liposomal hydrogels exhibiting a *ca*. 4-fold higher yield strain. To evaluate if lipid T_m_ impacted hydrogel mechanical properties, we evaluated hydrogels generated from low T_m_ (−17°C, dioleoyl tails) and high T_m_ (55°C, distearoyl tails) phospholipids (Supplemental Figure 5). Despite the significant difference in T_m_, these hydrogels exhibited consistent mechanical properties, except for the storage modulus G’, which was moderately elevated in the high T_m_ formulation. Overall, these data provide compelling evidence that liposomal hydrogels are robust to the liposome formulation. This versatility makes this platform broadly attractive for the conversion of any liposome-based nanotechnology into an injectable hydrogel.

### Lipid content and liposome size influence rheological properties of liposomal hydrogels

The mechanical properties of conventional hydrogels are tuned by changing the concentration of solids and crosslinkers in the system. Since the liposomal building blocks serve as crosslinkers in our hydrogel system, we set out to determine if lipid content could tune the mechanical properties of liposomal hydrogels. We found that as lipid content increased from 1 to 10 wt%, the resulting hydrogels became both stiffer and more solid-like across a broad range of frequencies (Figure 2A-B, D). For all formulations, no crossover was observed between the G’ and G’’ across the frequency range examined, indicating robust viscoelasticity and hydrogel formation across a wide range of lipid contents. Likewise, amplitude sweeps for each formulation demonstrated very high strains upon yielding and smooth yielding transitions with no brittle fracture.

**Figure 2.**
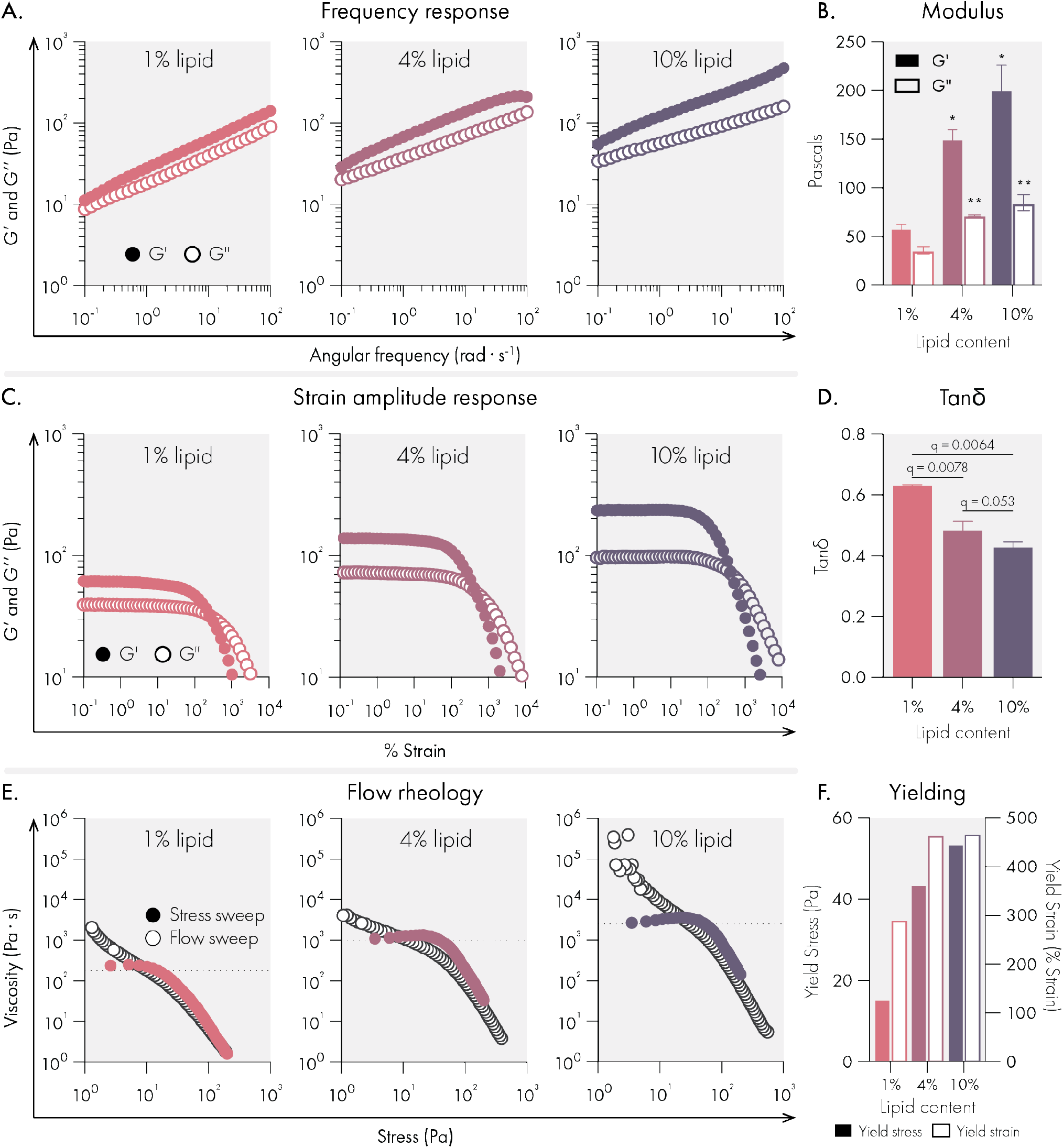
Mechanical properties of liposomal hydrogels are tuned by liposome concentration in the hydrogel. All measurements were performed on hydrogels composed of 1%, 4%, and 10% 50 nm extruded liposome content (by weight). All hydrogel formulations were 2% HPMC-C_12_ by weight. **(A)** Oscillatory shear rheology measurements indicate increased liposome content increases G’ and G’’. **(B)** G’ and G’’ values at 10 rad/s, data represent mean and SEM of 2 replicate batches. **(C)** Strain controlled oscillatory shear rheology. **(D)** Tan(delta) values at 10 rad/s, data represent mean and SEM of 2 replicate batches. **(E)** Flow rheology overlay showing low to high stress-sweep behavior (solid circles and high to low flow sweep (empty circles). **(F)** Yield stress and strain values extracted from flow rheology and strain amplitude response data. Statistical comparisons were made using a one-way ANOVA, and the false discovery rate (FDR) was controlled at 5% using the Benjamini, Krieger, and Yekutieli method.

While each formulation yielded a high-quality hydrogel, we observed that higher lipid content led to higher yield strains (Figure 2C). To characterize the yielding behavior of these materials more fully, we performed flow sweeps to assess each formulations’ yield stress behavior. Both shear rate-imposed flow sweeps (from high shear rate to low shear rate) and stress sweeps (from low stress to high stress) were performed (Figure 2E-F). These data indicated that all formulations exhibit extreme shear-thinning behavior, minimal thixotropy (as shown by the high degree of overlap in our two methods), and significant yield stress values (as indicated by the stress at which the viscosity abruptly decreases in the stress sweep). Similar to what we observed with yield strains, the yield stress of the hydrogels increased with higher lipid content. Overall, the high yield stress values noted for these formulations suggest that these materials are ideal for forming and maintaining subcutaneous depots robust to the low but constant stresses imposed by the dermis.

Next, the effect of liposome size on mechanical properties was investigated. As liposome size is increased at a constant mass loading in the hydrogel, there are fewer liposomes per unit volume. This decrease in the number of liposomes leads to greater interparticle spacing^35,36^ and reduces the total liposomal surface area with which the cellulose polymers can interact, likely reducing the efficiency of crosslinking interactions. Our data confirm these hypotheses and suggest that increasing liposome size leads to less stiff hydrogels with constant solid-like properties (Figure 3A-B, D). While stiffness (G’) and solid-like properties (tan delta = G’’/G’) are often correlated, it is fascinating that this modular system exhibits the ability to independently tune stiffness without changing the solid-like properties. In contrast, changing liposome size did not significantly affect the yield strain, though increases in yield stress were observed with the smallest liposomes (Figure 3, E-F).

**Figure 3.**
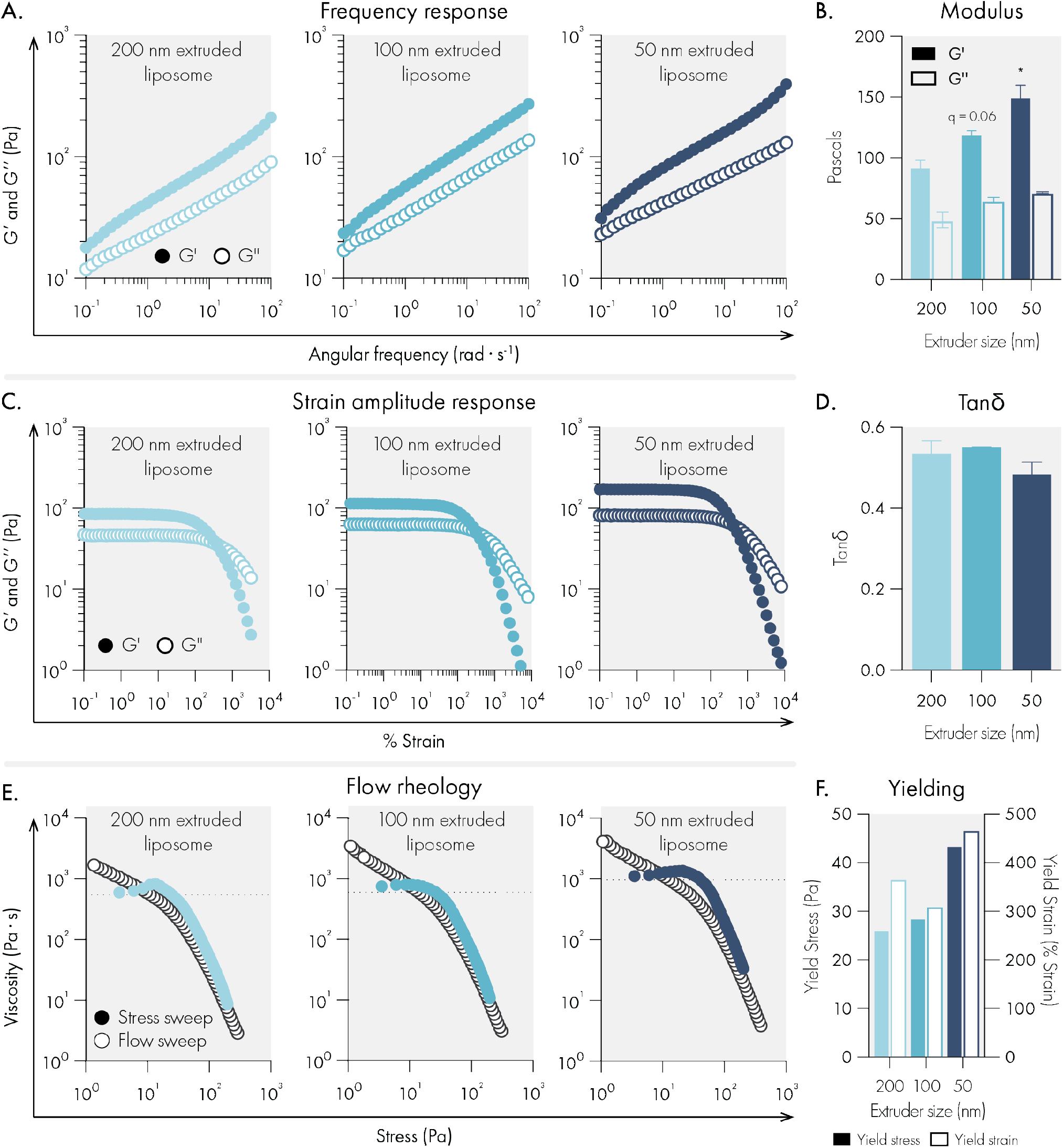
Liposome size impact on rheological properties of liposomal hydrogels. All measurements were performed with hydrogels composed of 2% HPMC-C_12_ and 4% liposome, by weight. Liposomes were extruded through either 200, 100, or 50 nm polycarbonate filters. **(A)** Representative oscillatory shear rheology. **(B)** G’ and G’’ at 10 rad/s indicates G’ is influenced by liposome size. Data represents the mean and SEM of two replicate batches. **(C)** Representative strain-controlled oscillatory shear rheology. **(D)** Tan(delta) of liposomal hydrogels is unchanged by liposome size. Data represents the mean and SEM of two replicate batches. **(E)** Representative flow rheology overlay showing low to high stress-ramp behavior (solid circles and high to low flow sweep (empty circles). **(F)** Yield stress and strain values extracted from flow rheology and strain amplitude response data. Statistical comparisons were made using a one-way ANOVA, and the false discovery rate (FDR) was controlled at 5% using the Benjamini, Krieger, and Yekutieli method.

While often neglected in the analysis of biomaterials, extensional rheology is highly important for material handling and processing in applications such as 3D printing and adhesives.^37^ For example, extensible materials have been demonstrated to improve 3D bioprinting fidelity.^38^ Upon handling our liposomal hydrogels, we noted both sticky and extensional characteristics. To further investigate these attributes, we performed filament stretching rheology to assess the formulations’ strain-to-break (Figure 4). Notably, we observed extreme extensibility for all formulations across several strain rates that are higher than all other physically crosslinked biomaterials reported to date.^37^ Intriguingly, we observed a moderate but significant increase in extensibility with an increase in liposome size (Figure 4D), potentially due to the increase in the interparticle spacing.

**Figure 4.**
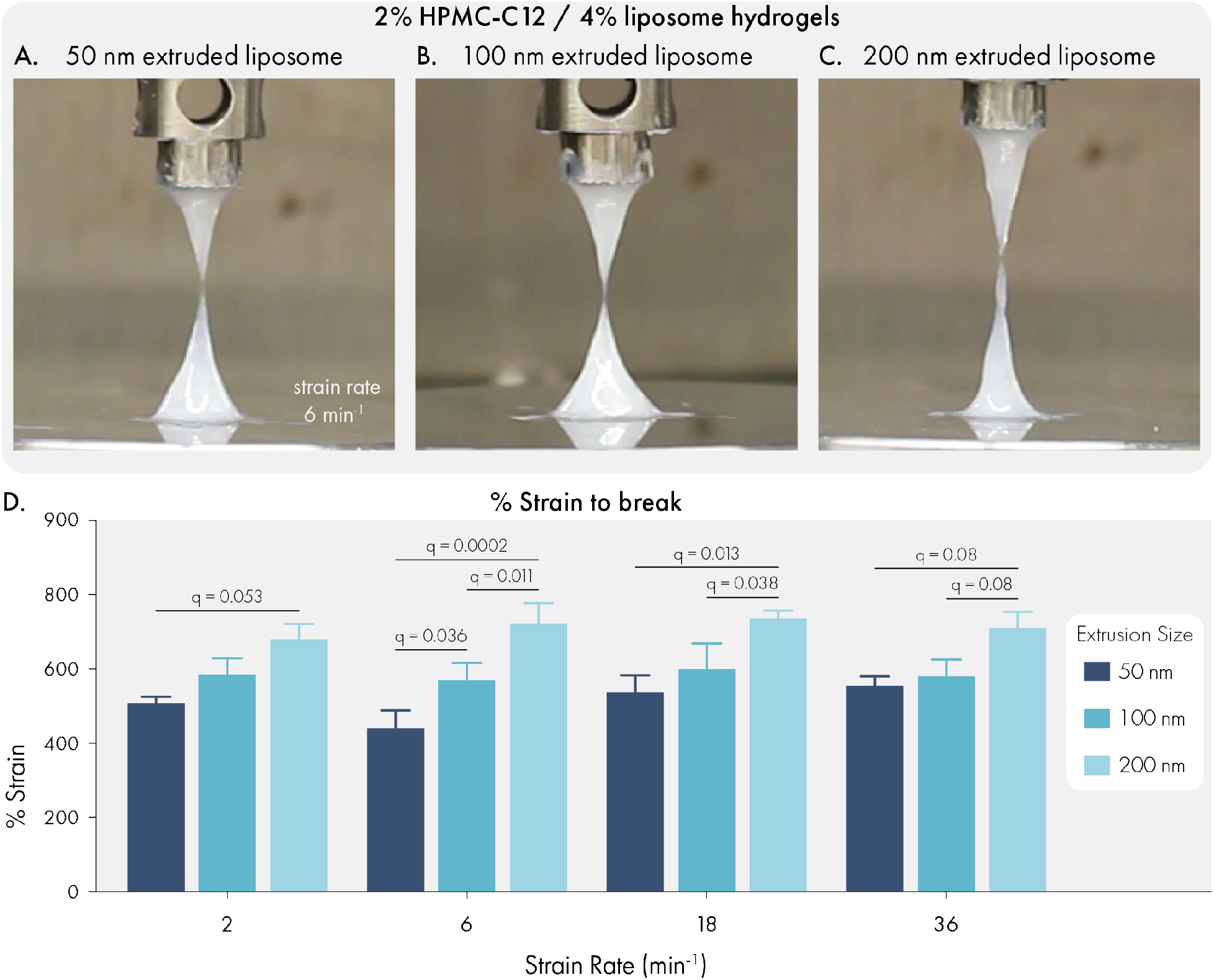
Liposome size influences the extensional properties of liposomal hydrogels. All measurements were performed with hydrogels composed of 2% HPMC-C_12_ and 4% liposome, by weight. Liposomes were extruded through either 200, 100, or 50 nm polycarbonate filter. Representative images of filament breakage with hydrogels composed of liposomes extruded through **(A)** 200 nm, **(B)** 100 nm, or **(C)** 50 nm polycarbonate filters. **(D)** Quantification of % strain to break for each formulation at a range of strain rates. Data represents mean and SEM of 3 independent batches. Statistical comparisons were made using a two-way ANOVA, and the false discovery rate (FDR) was controlled at 5% using the Benjamini, Krieger, and Yekutieli method.

### Liposomal hydrogels are non-toxic and recruit cell infiltrates in healthy mice

To confirm that liposomal hydrogel technology is suitable for biomedical applications, we conducted toxicological studies in immuno-competent C57BL6 mice. We injected 100 μL doses of liposomal hydrogel (2% HPMC-C_12_ polymer; 4% liposome; 2P-4L) subcutaneously into the hind flank of mice, where we observed the material take on a well-defined spherical shape (Figure 5A). Over the course of the following week, we monitored the mice daily for signs of skin irritation, weight loss, or any signs of morbidity, and we also tracked the size of the hydrogel depot using calipers. We did not observe any signs of skin irritation with the 2P-4L formulation (Figure 5B-C), and we noted that this formulation slowly dissolved over time until the material was explanted on day 7 (Figure 5D). Caliper measurements of the hydrogel area indicated that the depot was steadily eroding after implantation (Figure 5E), indicating that liposomal hydrogels are cleared without signs of toxicity or inflammation. This observation was further bolstered by the stable body weights of hydrogel-treated mice (Figure 5F), which exhibited consistent weights compared to untreated control animals.

**Figure 5.**
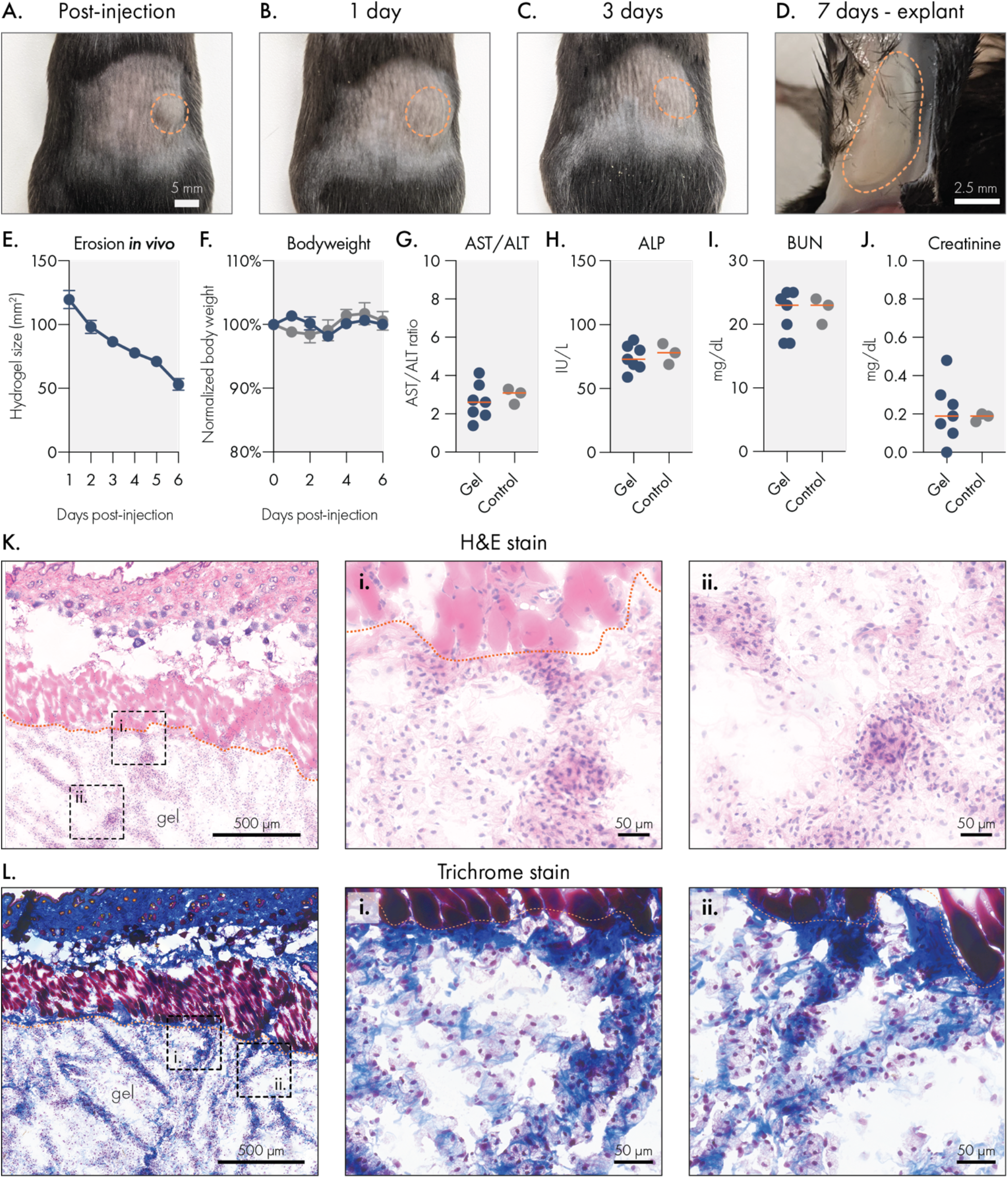
Liposomal hydrogels are biocompatible and biodegradable *in vivo*. Liposomal hydrogels were injected subcutaneously into the flanks of immuno-competent C57BL6 mice. Representative images of hydrogels **(A)** immediately after injection, **(B)** 1 day after injection, **(C)** 3 days after injection, and **(D)** upon explantation on day 7. Dotted orange line indicates edge of hydrogel depot. **(E)** Caliper measurements of hydrogel depot size over time indicates steady erosion of the material *in vivo*. **(F)** Stable body weights of hydrogel treated mice are consistent with body weights of untreated mice. Blood chemistry panel on serum collected 7 days after hydrogel injection. Liver function was tracked by measuring **(G)** AST/ALT ratio and **(H)** ALP levels. Kidney function was tracked by measuring serum **(I)** BUN and **(J)** creatinine levels. All blood chemistry measures were consistent between hydrogel treated and untreated control mice. Histological analysis of explanted liposomal hydrogels using **(K)** H&E or **(L)** trichrome staining.

Seven days after hydrogel injection, we collected serum from the mice to evaluate common markers of liver and renal toxicity. We observed that hydrogel-treated mice had comparable AST/ALT ratios and ALP levels to untreated control mice (Figure 5H-I), which indicates that these hydrogels do not induce acute hepatotoxicity after administration. Similarly, we found that hydrogel-treated and control mice had similar BUN and creatinine levels, suggesting that no short-term renal toxicity occurs after injection with liposomal hydrogels. Overall, these data indicate that liposomal hydrogels do not pose a risk of acute liver or kidney damage.

To determine how the body engages with liposomal hydrogels, we explanted the gels after 7 days and processed them for cryohistology (Figure 5K-L). H&E staining of the hydrogel sections revealed robust cellular infiltration into the biomaterial. Interestingly, cellular infiltrates are localized along strands of ECM that are deposited within the hydrogel. In these samples, there is little classical fibrosis along the hydrogel-host interface, and there are no signs of multi-nucleated foreign body giant cells. Overall, these observations indicate that this material does not evoke a strong foreign body response. Trichrome staining was then used to more closely evaluate the deposition of ECM in and around the hydrogels. This staining method, which renders collagen blue and muscle tissue red, revealed that the strands of tissue deposited into the hydrogel contain abundant collagen. In general, the toxicological and histological data indicate that liposomal hydrogels are well tolerated in the subcutaneous space and that these hydrogels are easily infiltrated by resident cells, and in particular macrophages. The ability for cells to infiltrate and remodel this biomaterial presents a promising opportunity for engineering specific cellular niches, such as those currently being explored by immuno-modulatory biomaterials.^39,40^

### Engineering the surface chemistry of liposomal building blocks enables distinct modes of release for protein cargo

Because protein-based therapeutics are becoming increasingly vital to diverse biomedical applications, we investigated strategies that would improve control over the release of proteins from liposomal hydrogels (Figure 6A). Initially, we investigated the effect of molecule size on diffusion within the hydrogel by conducting a Fluorescence-Recovery-After-Photobleaching (FRAP) experiment.^41^ In this study, we measured the mobility of fluorescein-labelled dextran molecules ranging from 40 to 2,000 kDa within the LNH system (Figure 6B). The data indicated that molecular weight greatly impacted diffusion (Figure 6C), confirming that the hydrogel mesh has the potential to slowly release high molecular weight (∼2,000 kDa) therapeutics, which may be useful for the delivery of bulky cargo (e.g., therapeutic biopolymers such as poly(I:C) adjuvants). In the 40 to 250 kDa size range (equivalent to a hydrodynamic radius between 4.5-12 nm, which also encompasses most protein therapeutics) we observed size-dependent release kinetics consistent with passive-release strategies from hydrogel networks.^7^ From these FRAP data, we estimate an apparent polymer mesh size of 4.45 nm.^42^

**Figure 6.**
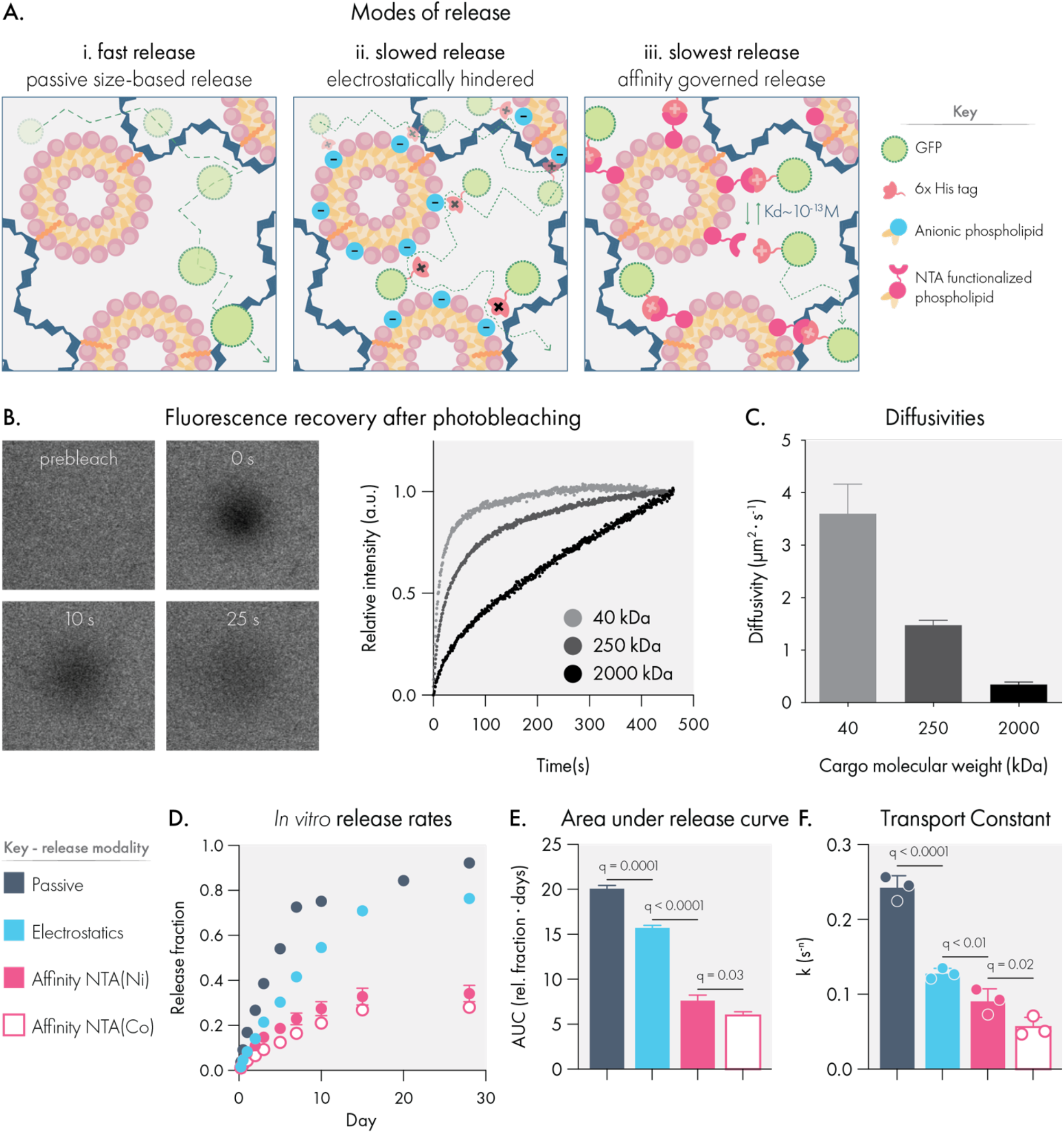
Liposome surface chemistry can be engineered to regulate the release rate of protein cargo. **(A)** Liposomal hydrogels feature three distinct release modalities based on cargo interaction with liposome surface chemistry. **(B)** Representative images and raw data for fluorescence recovery after photobleaching (FRAP) experiments. **(C)** FRAP data was used to calculate size-based diffusivity constants for model cargo (dextran macromolecules) within liposomal hydrogels. **(D)** In vitro release of his-tagged GFP from liposomal hydrogels engineered to engage the cargo through distinct mechanisms. Samples were maintained at 37°C throughout the study. **(E)** Area under the release time curve. **(F)** Release data were modeled using the Korsmayer-Peppas equation to calculate the transport constant. All data indicate mean and SEM of three independent replicate experiments. Statistical comparisons were made using a one-way ANOVA, and the false discovery rate (FDR) was controlled at 5% using the Benjamini, Krieger, and Yekutieli method.

We then explored two affinity-based strategies to regulate the release of protein drugs from these hydrogels by engineering the surface chemistries of our liposomal building blocks. First, we evaluated how electrostatic interactions influenced the release rate of a charged protein cargo (Figure 6A-ii). The DMPG phospholipid in our standard liposome formulation confers a net negative charge to these nanoparticles, which can interact with cationic amino acids. While electrostatic interactions have been leveraged in the past for delaying the release of charged proteins,^43-45^ these studies often rely on protein cargo that are intrinsically charged under physiological conditions. In this experiment, we instead introduced the common histidine-based tag (his-tag) onto proteins of interest. Because his-tags consist of a repeating stretch of cationic amino acids, we hypothesized that this motif would be sufficient to confer electrostatic interactions between his-tagged proteins and our anionic liposomal hydrogels.

Our *in vitro* release data confirmed that electrostatic engagement of the his-tag significantly reduced the release rate of GFP from the liposomal hydrogel. We observed that 50% of his-GFP cargo released from anionic liposomal hydrogels over 9 days, whereas 50% of wildtype GFP released from the same hydrogel matrix in only 4 days (Figure 6D). We attribute this to the inability of wildtype GFP to form electrostatic interactions with the anionic liposomes in this LNH formulation, whereas his-GFP is able to engage anionic liposomes through the repeating histidine motif. Over the course of the study, this led to significant differences (q = 0.001) in the area under the release time curve (Figure 6E), indicating his-GFP was better retained by the hydrogels. Fitting release data to the Korsmayer-Peppas model (Supplemental Figure 7),^46^ we estimated a significant 1.9-fold decrease (q < 0.0001) in the transport constant for his-GFP in the anionic liposomal hydrogel compared to wildtype GFP (Figure 6F). These data strongly indicate that engineering electrostatic interactions between the liposomes and protein cargo alters the release mechanism away from a passive diffusion-based mechanism to one governed by electrostatic affinity. Importantly, this effect was achieved by modifying protein cargo with the his-tag, a common epitope used in purification of recombinant proteins, which extends the utility of electrostatically governed release for many recombinant proteins.

We then set out to determine how specific affinity interactions would further regulate the release rate of protein cargo from this platform. We hypothesized that increasing the affinity for the his-tag motif would substantially slow the release of protein cargo from our material. To this end, we prepared liposomes featuring 3 mol % of either NTA(Ni) or NTA(Co)-functionalized phospholipids (Figure 6A-iii). These molecular motifs are commonly used in column purification of recombinant proteins, exhibiting extremely high affinity (K_d_ ∼ 10^−13^ M) for the his-tag motif.^47,48^ We observed that both NTA(Ni) and NTA(Co) functionalized liposomal hydrogels dramatically reduced the release rate of his-GFP *in vitro*, with only 34.1±0.1% and 28.0±0.1% of his-GFP released by 30 days, respectively. We noted a slower release rate from NTA(Co) formulations, possibly due to the partial oxidation of Co^2+^ to the Co^3+^ state during synthesis, which has been reported to increase the affinity to his-tag.^49,50^ The area under the release curve for NTA-based formulations were significantly reduced (q < 0.0001) compared to both passive and electrostatic release mechanisms. We also noted that NTA(Co) formulations reduced the AUC compared to NTA(Ni) formulations (q = 0.03).

Consistent with reports that NTA(Co) can exhibit higher affinity for the his-tag motif, the transport constant of his-GFP in NTA(Co) LNH formulations was 38.5% lower than that of NTA(Ni) (q = 0.02; Figure 6F). Impressively, the NTA(Co) affinity-governed release rate is 4.3-fold and 2.3-fold slower than the passive and electrostatic modes of release, respectively. These results are consistent with prior reports leveraging NTA-functionalized phospholipids for tethering proteins onto liposomal drugs, which observed stable loading of his-tagged proteins onto these constructs.^51^ While prior studies have focused on the utilization of functionalized liposomes alone for delivery,^52,53^ we report the first study to incorporate functionalized liposome as a structural component of an injectable hydrogel depot for applications in extended delivery.

Importantly, these data indicate that by engineering the surface chemistry of the liposomal building blocks of the LNH system distinct modes of release can be employed to deliver protein drugs on different timescales. Notably, LNHs release proteins over extremely long timescales compared to previously published reports.^54-56^ Of previously reported systems, physical networks functionalized with peptide binding domains show the most similar sustained release behavior.^57,58^ Moreover, as these strategies take advantage of common motifs (*e*.*g*., his-tag) used in the production of recombinant proteins, they also allow for orthogonal controlled release of multiple proteins based on the presence or absence of specific molecular handles.

### Liposomal hydrogels enable synchronized sustained release of antibodies and cytokine *in vivo*

Because liposomal hydrogels are able to simultaneously mediate orthogonal modes of release, we hypothesized that this biomaterial could be used to synchronize the release kinetics of different sizes and types of protein drugs. With conventional hydrogels, the release kinetics of multiple co-encapsulated proteins are often dictated by cargo size, with smaller proteins releasing more quickly than larger cargo.^7^ The inability to tune relative release rates between cargo is a major limitation for biomaterials, particularly for applications in immuno-engineering and tissue regeneration where the order of drug presentation can strongly influence outcomes.^3^

To initially test our hypothesis, we modeled hydrogel mediated co-delivery of IgG antibody and the IL-12 cytokine, two important classes of immunotherapy drugs. Because IgG antibodies are relatively bulky (*ca*. 150 kDa and estimated R_h_ of *ca*. 5.29 nm)^59^, we hypothesized that IgG would be effectively retained in our hydrogels using a passive-release mechanism. Using a previously published multiscale model for predicting passive release from hydrogel systems,^42^ we estimated the diffusivity of IgG to be 36.9 μm^2^/s in a conventional 10wt% PEG hydrogel.^60^ In contrast, we observed a much lower diffusivity (1.3 μm^2^/s) of IgG antibody in our LNH system, a roughly 30-fold decrease in IgG diffusivity.

Overall, our model predicted a highly effective sustained release of IgG from liposomal hydrogels. In contrast, IL-12 is a much smaller protein (*ca*. 60 kDa and estimated R_h_ of *ca*. 3.18 nm)^59^ that we anticipated would be poorly retained using passive release, aligning with our *in vitro* findings demonstrating rapid passive release of GFP (*ca*. 40 kDa). To begin evaluating this hypothesis, we modeled the diffusivity of IL-12 in our LNH system, which predicted that under passive release conditions, IL-12 would diffuse at a rate of 16.5 μm^2^/s, which is roughly 13-fold faster diffusivity than IgG. Taken together, our model suggests that purely passive release mechanisms would be incapable of synchronizing the sustained release of these two proteins from a drug delivery vehicle.

To synchronize the release kinetics of these distinctly sized proteins, we leveraged electrostatic interactions to slow the release of IL-12. Interestingly, IL-12 is capable of engaging with charged biomaterials for local drug delivery,^61^ and engages in electrostatic interactions with densely anionic heparin proteoglycans and endogenous components of the extracellular matrix.^62,63^ Given these properties, we hypothesized that our LNH system featuring anionic liposomes would be able to dramatically slow the release of IL-12, similar to our *in vitro* results.

To test this hypothesis, we subcutaneously co-delivered fluorescently labelled IgG and IL-12 proteins in liposomal hydrogels or bolus injections in immunocompetent SKH1E mice (Figure 7A) and analyzed the release of both proteins simultaneously with an *in vivo* imaging system (IVIS). Each protein was labelled with a unique non-overlapping fluorophore to enable simultaneous tracking of both proteins. IVIS imaging revealed that bolus administration led to a rapid decline of both IgG and IL-12 in less than one day. In contrast, liposomal hydrogels closely aligned the release profiles of these two very differently sized proteins. Even after 14 days, the liposomal hydrogels retained over 50% of the fluorescent signal from both proteins.

**Figure 7.**
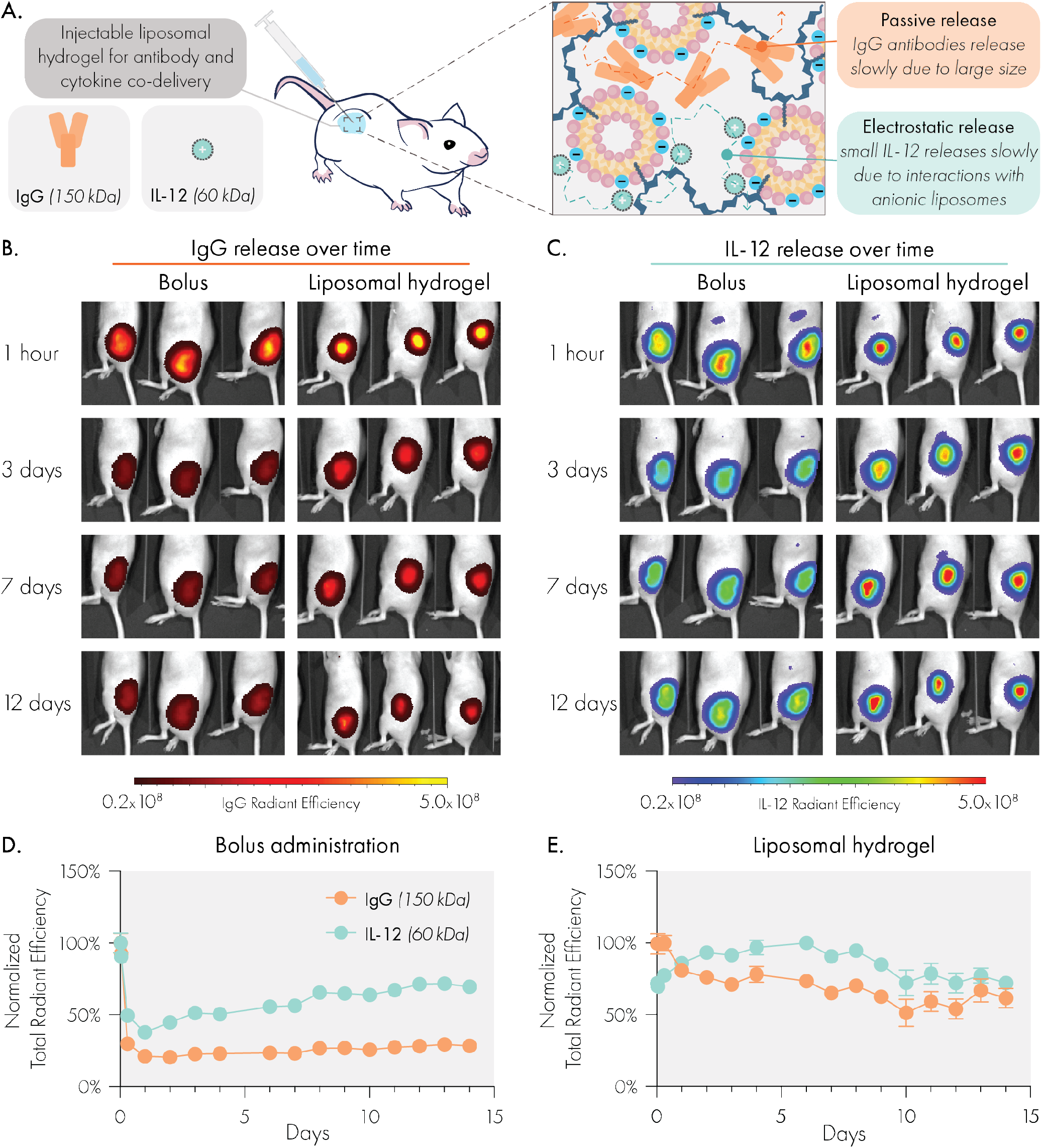
Liposomal hydrogels enable synchronized and sustained co-delivery of IgG antibody and IL-12 cytokine cargo *in vivo*, despite substantial size difference. **(A)** Liposomal hydrogel mesh size is able to significantly slow release of bulky IgG antibodies, while anionic liposomes regulate release of small IL-12 cytokines via electrostatic interactions. Representative IVIS images of **(B)** Alexa Fluor 594 labeled IgG and **(C)** Alexa Fluor 700 labeled IL-12 release over time. Quantified release curves of IgG and IL-12 following **(D)** local subcutaneous bolus injection or **(E)** subcutaneous injection of liposomal hydrogel. Sample size was n=3 for bolus and n=4 for hydrogel groups.

We note that in the bolus injections, the release curves plateau, indicating some fraction of cargo is retained in the injection site. We attribute this to immune cells in the injection site binding the cargo. While we chose isotype IgG and human IL-12 to reduce immuno-activity and simplify pharmacokinetics within this murine model,^64^ isotype antibodies still engage Fc receptors on myeloid cells and human IL-12 can still bind to local extracellular matrix constituents.^63,65^ Similar to our prior fluorescence-based pharmacokinetic studies in hydrogels,^66^ we noted a self-quenching phenomenon affecting the IL-12 signal from liposomal hydrogels over our early timepoints. During this phase, signal increases as fluorescent molecules separate from one another and cease self-quenching.

This study demonstrates the unique capabilities of LNHs to program release of highly physicochemically distinct and translationally relevant molecular cargo *in vivo*. Previously published affinity-based release systems have primarily demonstrated efficacy *in vitro*.^*54,55,58*^ This study is among the first to demonstrate programmable release over uniquely long timescales in a fully immunocompetent *in vivo* setting.

## Conclusion

We have developed a novel injectable liposomal nanocomposite hydrogel biomaterial and demonstrated its exceptional utility in enabling programmable release of diverse protein cargo both *in vitro* and *in vivo*. Comprehensive shear and extensional rheology demonstrate that liposomal hydrogels are highly tunable and modular materials which exhibit gel-like properties, facile injectability, and rapid self-healing ideal for forming drug-releasing depots following administration. Our study reveals that hydrogel formation is highly robust to modifications of the liposomal building blocks, especially with regards to their surface chemistry. Additionally, the unique extensibility of these materials showcase promise for applications beyond drug delivery, such as 3D bioprinting or bioadhesives.

To highlight the translational potential of this material, we investigated its biodegradability and biocompatibility in mice through body weight tracking, blood chemistry, and histology. Overall, LNHs eroded gradually *in vivo* and caused no adverse side effects. We also demonstrated multi-modal release mechanisms from the LNH platform that ranged from passive to affinity-based release. FRAP studies indicated that liposomal hydrogels regulate diffusion of bulky protein cargo passively through their small dynamic mesh size. For cargo smaller than the apparent mesh size, we demonstrated two modes of affinity-governed release based on either electrostatic interactions or affinity ligands. In the case of electrostatics, charged liposomes within the LNH system engaged oppositely charged protein cargo. In the case of affinity ligands, NTA-functionalized liposomes within the LNH system selectively bound to his-tagged proteins. In both cases, the release rate of small proteins was reduced to various extents to provide desired release kinetics. Finally, we demonstrated this material’s ability to synchronize the delivery of IgG and IL-12, two clinically relevant and differently sized biomolecules, *in vivo*. We found that liposomal hydrogels enabled weeks of sustained, synchronized, and local release of these two distinct protein drugs.

Overall, this report highlights an approach leveraging liposomal nanotechnology to generate a modular, safe, and effective biomaterials platform for coordinating the release kinetics of multiple protein drugs. The ability to inject this material further increases the clinical and translational potential of this technology. These advanced capabilities have impactful implications for a variety of biomedical applications requiring the controlled delivery of multiple drugs, particularly in the areas of tissue and immuno-engineering.

## Materials and Methods

### Materials

All lipids were purchased from Avanti Polar Lipids. See Supplemental Table 1 for a complete list of lipids and their associated catalog numbers. Hypromellose, N-methylpyrrolidone (NMP), and dodecyl isocyanate were purchased from Sigma Aldrich. Dialysis tubing was purchased from Spectrum Labs. Sterile PBS was purchased from Thermo Fisher. Tag-free and histidine tagged GFP were purchased from MilliporeSigma and Sino Biologicals, respectively. Mouse IgG1 isotype control antibody (clone MOPC-21) was purchased from Bio X Cell. Tag-free and histidine tagged human IL-12 were both purchased from Sino Biologicals.

### Liposome synthesis

Liposomes were prepared via thin-film hydration and extrusion methods. Briefly, the desired amount of phospholipid was dissolved in chloroform or methanol (depending on the solubility of the particular phospholipid) and transferred to a round bottom flask. The flask was then attached to a Rotary Evaporator (Buchi) and placed under vacuum at 40°C until a thin lipid film was formed. The film was rehydrated in sterile PBS at a concentration ranging from 60 to 150 mg/mL. Rehydration occurred under rotary agitation and at 40°C for 30 minutes. Following rehydration, the crude liposomal solution was manually extruded through 400, 200, 100, and 50 nm polycarbonate filters (10 passes per each filter) using a hand-extruding device (either Avanti’s Mini Extruder or T&T Scientific’s NanoSizer MINI system). Size and uniformity of the liposomes were confirmed by dynamic light scattering measurements using the Wyatt DynaPro instrument. Liposomes were used immediately or stored at 4ºC until used.

### Dodecyl-modified hydroxypropylmethylcellulose (HPMC-C_12_) synthesis

HPMC-C_12_ was prepared as described previously.^24^ Briefly, 1 g of Hypromellose was dissolved in 40 mL of NMP and heated to 80 ºC in a PEG bath, with stirring. Dodecyl isocyanate (125 μL) was diluted in 5 mL of NMP and then added dropwise to the Hypromellose solution. HUNIGS catalyst was added dropwise (10 drops) to the mixture, after which the heat was shut off and the reaction allowed to continue overnight while stirring. The next day, polymer was precipitated in a 600 mL bath of acetone and then dissolved in 40 mL of Millipore water. The dissolved polymer was then purified via dialysis (3.5 kDa MWCO) over 4 days at room temperature. Pure HPMC-C_12_ was then lyophilized and dissolved in 1x sterile PBS to yield a 6wt% solution, which was stored at 4ºC until used.

### Liposomal hydrogel formation

Hydrogels were formed by simple mixture of liposomes and HPMC-C_12_ either in an Eppendorf tube or using a dual-syringe mixer. In an Eppendorf tube, the desired amount of liposome (measured by volume) and HPMC-C_12_ (measured by weight) were added, with any remaining volume adjustments being made by addition of sterile PBS. A metal spatula was then used to manually mix the solutions until a hydrogel was formed (2-3 minutes). For preparation using the dual-syringe method,^67^ the desired amount of liposome and HPMC-C_12_ solution were loaded into two separate 1 mL luer-lock syringes. The two syringes were then securely connected to one another via a luer-lock elbow connector. The solutions were mixed by alternating depression on the connected syringes until the hydrogel formed (determined by an increase in resistance during plunger depression, generally occurring after 100-150 passes). Hydrogels formulations generally consisted of 2wt% HPMC-C_12_ and varying weight percentages of liposome (either 1%, 4%, or 10%), depending on the particular experimental goals. Formulation shorthand is xP-yL where x indicates the weight percentage of HPMC-C_12_ and y indicates the weight percentage of liposome. Unless otherwise noted, liposome formulations are 9:1:2 molar ratios of DMPC, DMPG and cholesterol, extruded through 50 nm polycarbonate filters as described above.

### Rheological characterization of liposomal hydrogels

#### Shear Rheology

Rheological testing was performed using a 20 mm diameter serrated parallel plate at a 500 um gap on a stress-controlled TA Instruments DHR-2 rheometer, unless otherwise specified. All experiments were performed at 25 °C. Frequency sweeps were performed at a strain of 1% within the linear viscoelastic regime. Amplitude sweeps were performed at frequency of 10 rad/s. Flow sweeps were performed from high to low shear rates with steady state sensing. Stress sweeps were performed from low to high with steady state sensing

#### Viscometry

A Rheosense m-VROC viscometer was used to measure the hydrogel viscosity at high shear rates from low to high shear rates using a 1 mL Hamilton syringe. Each data point was collected at steady state.

#### Extensional Rheology

Strain-to-break measurements were performed on a TA Instruments ARES-G2 rheometer in axial mode with an 8 mm serrated plate geometry and Hencky strain rates as described by Nelson et al.^37^ A serrated parallel-plate with a radius of *R*_0_ = 4 mm and advanced Peltier system bottom plate were used. Samples containing 400 uL were loaded at a gap of *H*_0_ = 4 mm, resulting in an aspect ratio of Λ_0_ = *H*_0_/*R*_0_ = 1. The serrated plate helped to ensure the material would cohesively fail. All experiments were performed at 25 °C and replicated three times from independent batches of hydrogel.

### Toxicology studies

All animal studies were performed following Stanford’s IACUC guidelines and protocols. Biocompatibility was assessed by tracking hydrogel erosion, body weight, and blood chemistry in mice subcutaneously injected with 100 μL of liposomal hydrogel. Mice were injected with hydrogels subcutaneously in the hind flank, and the hydrogel size and mouse weight were tracked daily until the end of the study. During this time, the overall health of the mouse was also evaluated, in particular taking note of any skin irritation or signs of pain and distress (e.g., hunched posture or discontinuation of grooming behaviors). At the end of the study, mice were euthanized and blood collected via cardiac puncture. Blood was processed to recover serum and then submitted to Stanford’s Animal Diagnostic Lab for blood work. Hydrogels were also explanted from mice and cryopreserved in OCT compound (Tissue Tek) using liquid nitrogen. Frozen specimens were then submitted to Stanford’s Animal Histology Services core facility for sectioning and staining. Stained slides were imaged using a Leica Thunder microscope using a 40x air objective.

### Fluorescence recovery after photobleaching experiments

Hydrogels were placed onto glass slides and imaged using a confocal LSM780 microscope at a 20x magnification. Samples were imaged using a low intensity laser to observe an initial level of fluorescence. Then the laser was switched to full intensity and focused on a region of interest (ROI) with a 25 um diameter for 10 seconds in order to bleach a circular area. Fluorescence data were then recorded for 4 minutes to create an exponential fluorescence recovery curve. Samples were taken from different regions of each gel (n=3). The diffusion coefficient was calculated according to,

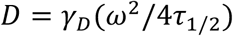

where the constant *γ*_D_ = *τ* _½_/*τ*_D_, with *τ* _½_ being the half-time of the recovery, *τ*_D_ the characteristic diffusion time, both yielded by the ZEN software, and *ω* the radius of the bleached ROI (12.5 μm).^41^

### *In vitro* release studies

Release of GFP from liposomal hydrogels was assessed by plate-reader analysis of serially collected samples as described previously. Briefly, 2P-4L hydrogels were prepared and used to encapsulate 10 μg of recombinant GFP protein (either tag-free or his-tag, depending on the experimental condition). To prepare specimens, approximately 100 μL of hydrogel was injected into the bottom of sealed glass capillary tubes. The remaining volume (200-300 μL) was filled with sterile PBS and the specimens were then incubated at 37°C in the dark. At the indicated time points, the supernatant was collected and replaced with fresh PBS. The collected supernatants were transferred to a black 96 well plate and analyzed using a plate reader. Three specimens were prepared for each experimental condition.

### Fluorescent labeling of proteins

Mouse isotype IgG (BioXCell; BE0083, clone MOPC-21) and human IL-12 (Sino Biological; CT011-H08H) were covalently modified with alexa fluor dyes using Abcam Lightning Link protein labeling kits according to the manufacturer’s instructions. IgG was labeled with Alexa Fluor 598 while IL-12 was labeled with Alexa Fluor 700, two non-overlapping dyes which would enable simultaneous tracking of both cargo with an IVIS imaging device. After labelling, both proteins were washed and concentrated using VivaSpin 500 desalting columns. Proteins were then mixed with liposomes prior to formation of hydrogels or adjusted to the desired concentration with sterile PBS for bolus administrations.

### *In vivo* release studies

SKH1E nude immunocompetent mice were used to assess the release behavior of LNHs containing labeled IgG and IL-12. Mice were subcutaneously injected in the hind flank with a 100 μL volume of either a bolus solution or a liposomal hydrogel. Each injection administered 10 μg of mouse IgG antibody and 12.5 μg of human IL-12. Isotype control IgG and human IL-12 were chosen for this study because both cargo were expected to have minimal immunoactivity, which would simplify our experimental system. Mice were imaged sequentially using a PerkinElmer IVIS instrument using automatic exposure acquisition settings. Mice were imaged immediately before injection, 30 minutes after injection, 1 hour after injection, 7 hours after injection, and then daily thereafter. Data were analyzed in the LivingImage software. Analysis consisted of defining a region of interest over the injection site or hydrogel, and exporting the total radiant efficiency measured in that region. Data were normalized to the maximum signal recorded during the study and then plotted in PRISM. Sample was n=3 for the bolus condition and n=4 for the liposomal hydrogel condition.

### Modeling

To model diffusivity of the IgG and IL-12, the hydrodynamic radius of IgG antibody and IL-12 were assumed to be 5.29 nm and 3.18 nm, respectively.^59^ The dynamic mesh size of the PLNP was approximated as 4.452 nm, back-calculated using our FRAP diffusivity measurements of dextran molecules of known hydrodynamic radii and utilizing the multiscale diffusion model for solute diffusion in hydrogels calculator derived in Axpe et al.^42,68^ Diffusivities of IgG and IL-12 were additionally calculated by the same method in standard PEG hydrogels for comparison by assuming a mesh size of 25 nm and a solid content of 10 wt%.^42^

### Statistical methods

Statistical comparisons were made in the GraphPad PRISM statistical software using either a one-way or two-way ANOVA, depending on the nature of the dataset. For all analyses, the false discovery rate was controlled at 5% for multiple comparisons using the Benjamini, Krieger, and Yekutieli method.

## Acknowledgements

The authors would like to thank the Soft Materials Facility, Animal Histology Services, and the Animal Diagnostics Lab at Stanford. We would also like to thank Professor Matthew Webber for helpful discussion of histology.

## Funding

This research was financially supported by the American Cancer Society (RSG-18-133-01-CDD), the Goldman Sachs Foundation (administered by the Stanford Cancer Institute, SPO# 162509), and a Bio-X Interdisciplinary Initiatives Seed Grant. S.C. is supported by the National Cancer Institute of the National Institutes of Health under Award Number F32CA247352. A.K.G. is thankful for a National Science Foundation Graduate Research Fellowship and the Gabilan Fellowship of the Stanford Graduate Fellowship in Science and Engineering.

## Author contributions

S.C. developed the initial concept for the material. S.C. and A.K.G. designed all experiments and analyzed the data. S.C., A.K.G., and J.H.K. carried out all the experiments. H.L.H. assisted with design of rheological experiments and data interpretation. S.C., A.K.G., and E.A.A. interpreted the results and wrote the manuscript.

## Competing Interests

S.C., A.K.G., J.H.K., and E.A.A. are listed as inventors on a patent application reporting the liposomal hydrogel technology described in this article (U.S. Patent Application No. 63/177373).

## Notes

### Competing Interest Statement

S.C., A.K.G., J.H.K., and E.A.A. are on a pending patent reporting the liposomal hydrogel technology described in this article (U.S. Patent Application No. 63/177373.

