## Supplemental Materials for "Injectable Liposome-based Supramolecular Hydrogels for the Programmable Release of Multiple Protein Drugs"

**Table of Contents**

Supplemental Table 1. Phospholipids used in study……………………………………………2
Supplemental Table 2. Liposome formulations..……………………………………………..3
Supplemental Table 3. Representative DLS for liposomes..………………………………...4
Supplemental Figure 1. Liposomes and HPMC-$C_{12}$ are required for gelation……………......5
Supplemental Figure 2. Rheological properties of cationic liposomal hydrogels………......6
Supplemental Figure 6. Effect of NTA-functionalization on liposomal hydrogels………...10
Supplemental Figure 7. Korsemeyer-Peppas fit of in vitro release data…………………..11
Supplemental Figure 8. Estimated diffusivity of IgG and IL-12 in hydrogels……………...12
Supplemental References ...……………………………………………………………………13

Supplemental Table 1. Phospholipids used to prepare liposomes in this study, and their physicochemical characteristics. All products were obtained from Avanti Polar Lipids.

| Name | Catalog Number | Molecular Weight (g/mol) | Net Charge (0=Neutral; +1=Positive; -1=Negative) | Melting Temperature (ºC ) |
|---|---|---|---|---|
| DSPC | 850365C | 790.145 | 0 | 55 |
| DSPG | 840465P | 801.058 | -1 | 55 |
| Cholesterol | 700100P | 386.65 | 0 | 0 |
| 18:1 DOPG | 840475P | 797.026 | -1 | -18 |
| 18:1 DOPC | 850375C | 786.113 | 0 | -17 |
| 14:0 DMPG | 850345C | 677.95 | -1 | 23 |
| 14:0 DMPC | 850345C | 677.95 | 0 | 24 |
| 18:1 DGS-NTA(Ni) | 790404 | 1057.003 | 0 | 0 |
| 18:1 DGS-NTA(Co) | 791113 | 1057.24 | 0 | 0 |
| DOPE | 850725C | 744.034 | 1 | -16 |
| DOPE-PEG(2000)-Cy5 | 880153C-1mg | 3234.11 | 1 | -16 |
| POPG | 840457C | 770.989 | -1 | -2 |
| DMG-PEG2000 | 880151 | 2509.2 | 0 | Not provided |
| 18:1 TAP (DOTAP) | 890890 | 698.542 | 1 | 0 |

Supplemental Table 2. Liposome formulations reported as mole ratios.

| Formulation | Lipids | Mole Ratio | Where Shown |
|---|---|---|---|
| 1 (Standard) | DMPC – DMPG – cholesterol | 9 – 1 – 2 | Figures 1, 2, 3, 4, 5, 6, 7 & Supplemental Figures 1, 4 |
| 2 (Medium cholesterol) | DMPC – DMPG – cholesterol | 9 – 1 – 6 | Supplemental Figure 4 |
| 3 (High cholesterol) | DMPC – DMPG – cholesterol | 9 – 1 – 10 | Supplemental Figure 4 |
| 4 (Cationic) | DMPC – DOTAP – cholesterol | 9 – 1 – 2 | Supplemental Figure 2 |
| 5 (PEGylated) | DMPC – DMPG – cholesterol | 13 – 2 – 4 – 1 | Supplemental Figure 3 |
| 6 (Low Tm) | DOPC – DOPG – cholesterol | 9 – 1 – 2 | Supplemental Figure 5 |
| 7 (High Tm) | DSPC – DSPG – cholesterol | 9 – 1 – 2 | Supplemental Figure 5 |
| 8 (NTA-Ni functionalized) | DMPC – DMPG – cholesterol – DGS-NTA(Ni) | 9 – 1 – 2 – 0.379 | Figure 6 & Supplemental Figure 6 |
| 9 (NTA-Co functionalized) | DMPC – DMPG – cholesterol – DGS-NTA(Co) | 9 – 1 – 2 – 0.379 | Figure 6 |

Supplemental Table 3. Representative Dynamic Light Scattering of 50 nm extruded liposomes. Several Acquisitions of the Same Sample.

| Item | Diameter (nm) | Normalized Intensity (Cnt/s) | Radius (nm) | %PD | Mw-R (kDa) | PD Index | Polydispersity (nm) |
|---|---|---|---|---|---|---|---|
| Acq 1 | 91.4 | 46655701 | 45.7 | 13 | 25769 | 1.31E-01 | 6 |
| Acq 2 | 91.5 | 47009476 | 45.77 | 8.1 | 25859 | 8.07E-02 | 3.7 |
| Acq 3 | 89.4 | 45015109 | 44.72 | 10.1 | 24497 | 1.01E-01 | 4.5 |
| Acq 4 | 89.7 | 46480861 | 44.87 | 15.2 | 24686 | 1.52E-01 | 6.8 |
| Acq 5 | 91.6 | 45287323 | 45.78 | 4.2 | 25867 | 4.17E-02 | 1.9 |
| Acq 6 | 91.1 | 46959625 | 45.56 | 4.5 | 25586 | 4.49E-02 | 2 |
| Acq 7 | 89.2 | 46312122 | 44.61 | 12.5 | 24355 | 1.25E-01 | 5.6 |
| Acq 8 | 90 | 45467863 | 45.02 | 8.9 | 24881 | 8.92E-02 | 4 |
| Acq 9 | 91.6 | 47301599 | 45.81 | 13.6 | 25906 | 1.36E-01 | 6.2 |
| Acq 10 | 88.9 | 45506154 | 44.46 | 11 | 24164 | 1.10E-01 | 4.9 |

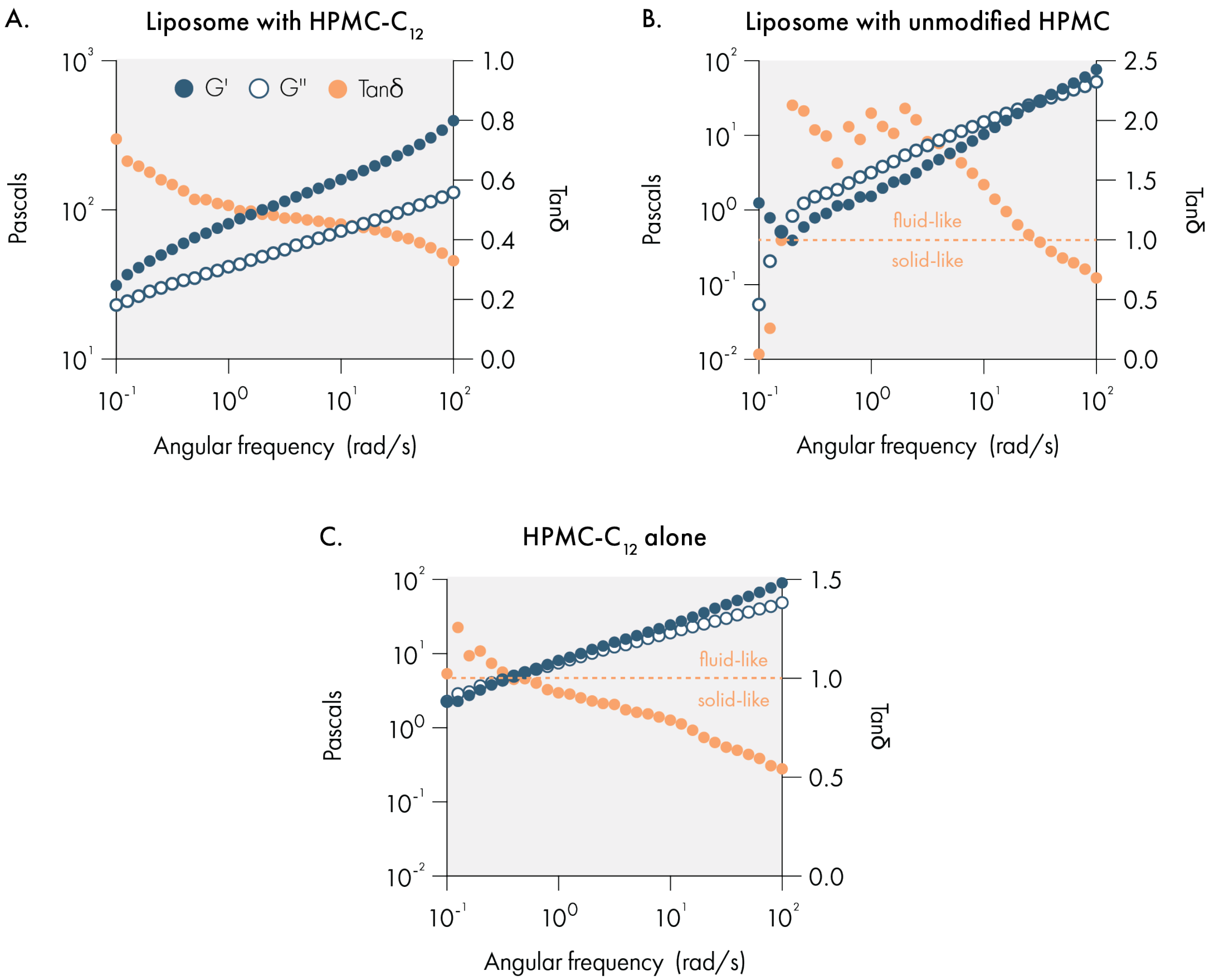

**Supplemental Figure 1.** Frequency sweep of liposomal hydrogels incorporating 2wt% hydroxypropylmethylcellulose (HPMC) with (**A**) and without (**B**) hydrophobic dodecyl ($C_{12}$) modification. Both formulations were prepared with 10wt% liposome content. **(C)** Rheological characterization of a 2wt% solution of HPMC- $C_{12}$ without any liposomes. Elastic storage modulus (G') and viscous loss modulus (G'') across a wide range of frequencies on left y-axis. Tan(*delta*) is shown on the right y-axis, which can be used to determine whether the material exhibits solid or fluid properties. Dotted line indicates which Tan(*delta*) values indicate solid and liquid states.

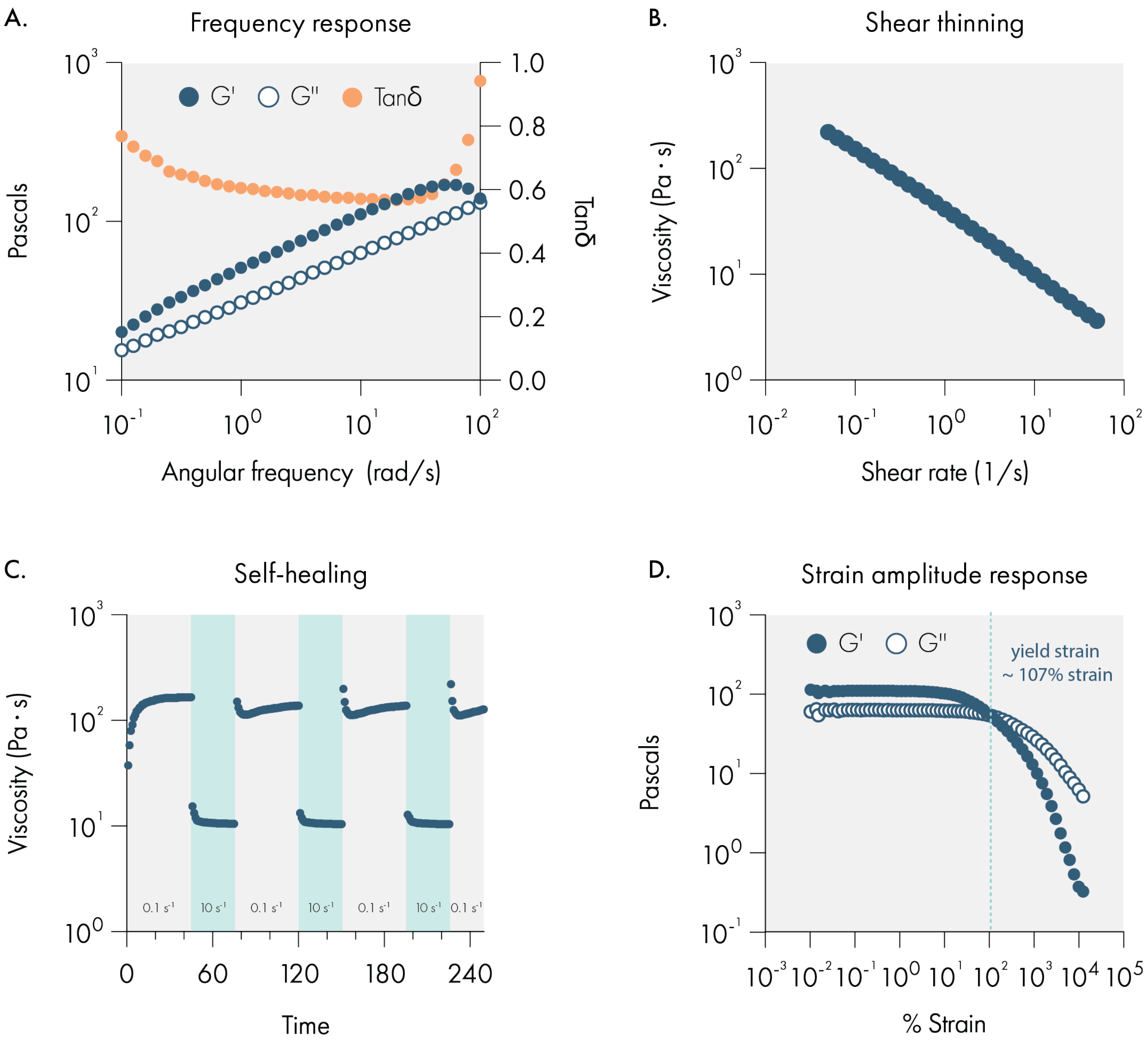

**Supplemental Figure 2.** Rheological properties of cationic liposomal hydrogels. (**A**) Frequency sweep demonstrating a gel-like response. (**B**) Flow sweep demonstrating the shear-thinning behavior. **(C)** Self-healing behavior demonstrated by cycling between a high (10 $s^{-1}$) and low (0.1 $s^{-1}$) shear rates with rapid recovery of the viscosity upon reduction of the shear rate. **(D)** Amplitude sweep assessing the linear viscoelastic regime and demonstrating the high % strain at yielding.

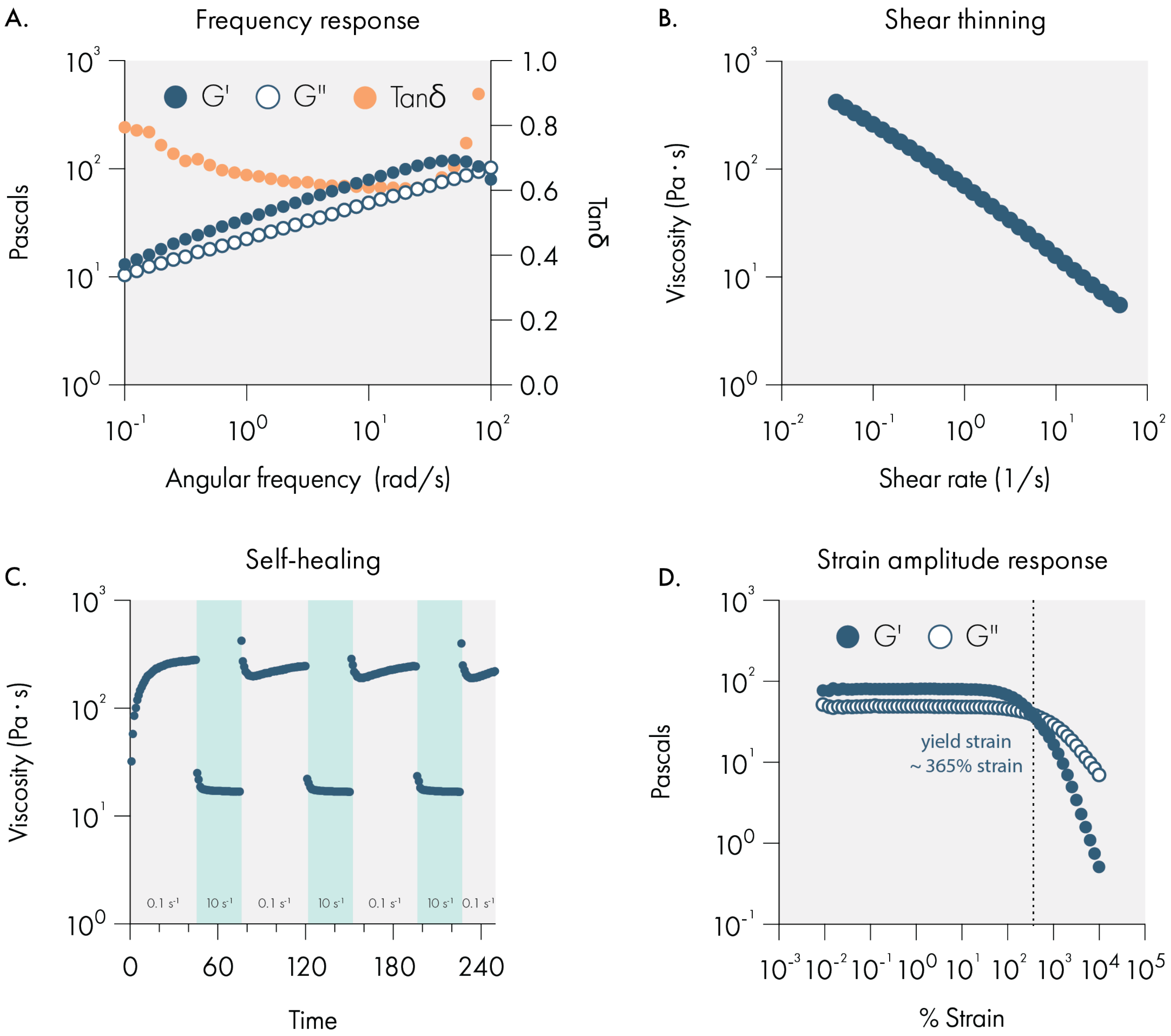

**Supplemental Figure 3.** Rheological properties of PEGylated liposomal hydrogels. **(A)** Frequency sweep demonstrating a gel-like response. **(B)** Flow sweep demonstrating the shear-thinning behavior. **(C)** Self-healing behavior demonstrated by cycling between a high (10 s-1) and low (0.1 s-1) shear rates with rapid recovery of the viscosity upon reduction of the shear rate. **(D)** Amplitude sweep assessing the linear viscoelastic regime and demonstrating the high % strain at yielding.

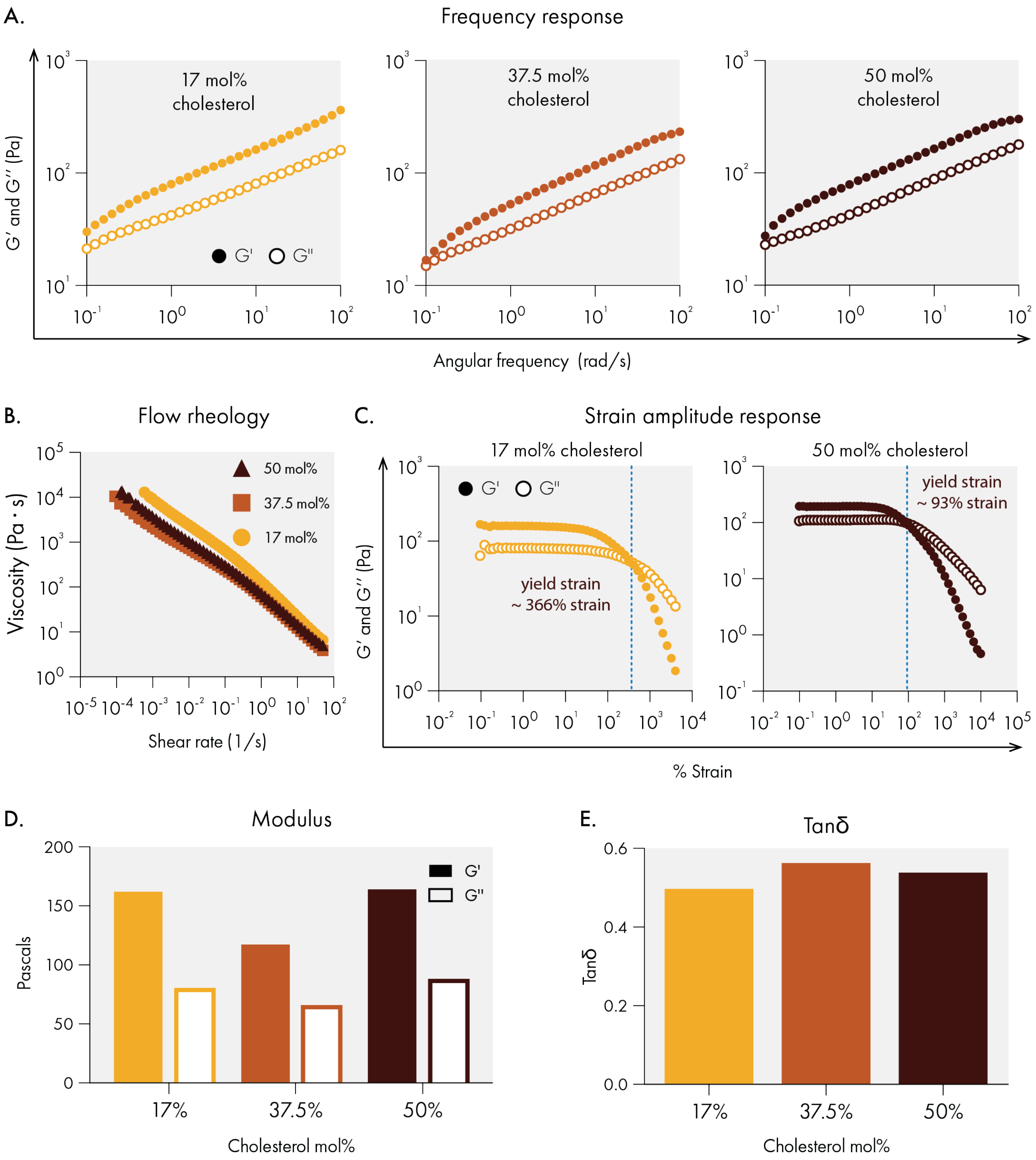

**Supplemental Figure 4.** Effect of cholesterol content on liposomal hydrogel rheological properties. **(A)** Frequency sweep demonstrating the relatively consistent gel-like response across several cholesterol mol% additions. **(B)** Flow sweep demonstrating the relatively consistent shear-thinning behavior across several cholesterol mol% additions. **(C)** Amplitude sweep assessing the linear viscoelastic regime and demonstrating the high % strain at yielding across several cholesterol mol % additions. **(D)** Elastic storage moduli (G') and viscous loss moduli (G'') at 1% strain and 10 rad/s of several cholesterol mol% additions. **E.** Tan(delta) at 1% strain and 10 rad/s of several cholesterol mol% additions.

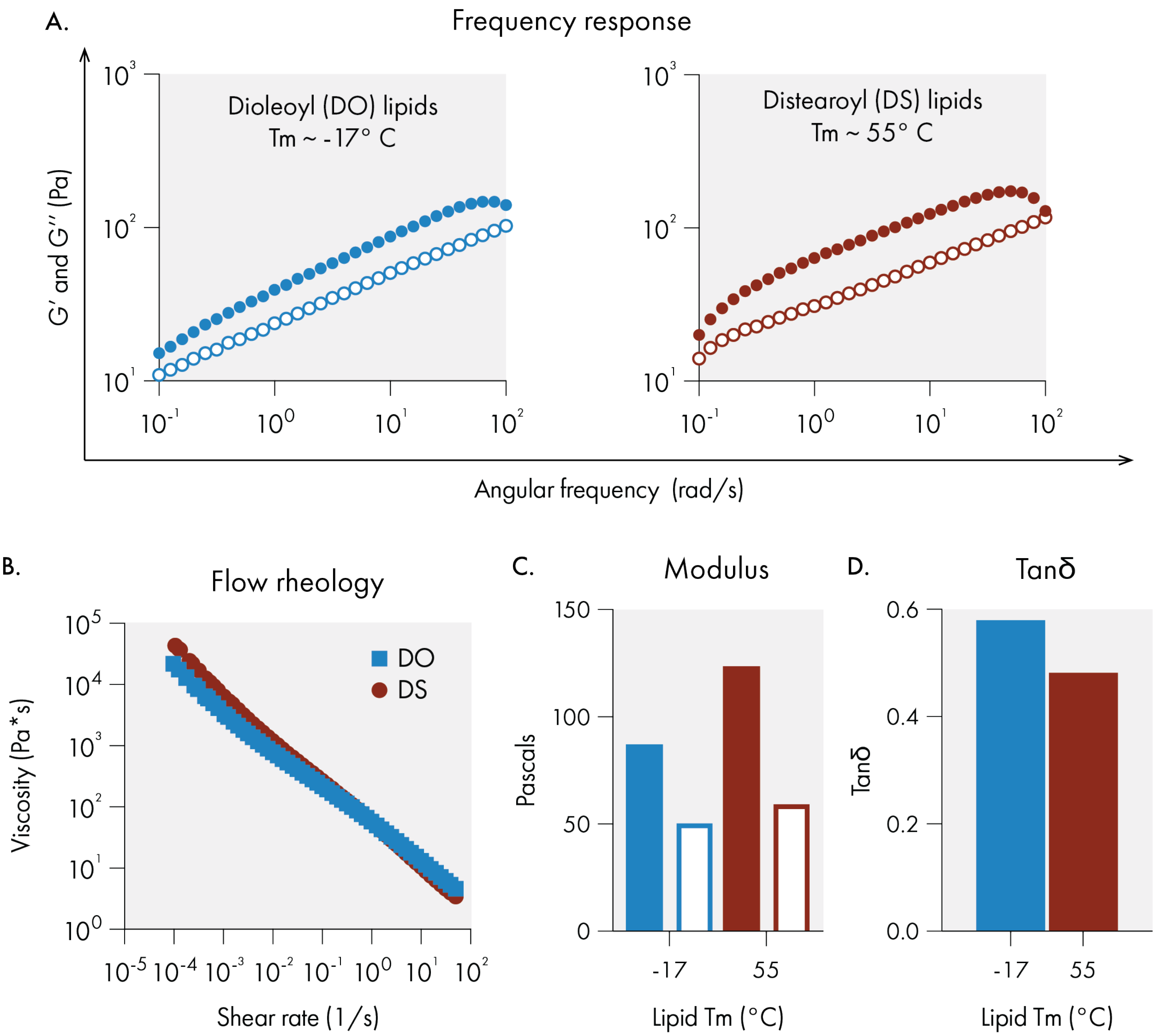

**Supplemental Figure 5.** Effect of varying the melting temperature (Tm) of the lipids comprising the liposomes in liposomal hydrogels. Dioleoyl lipids were incorporated for a low Tm, Distearoyl lipids were incorporated for a high Tm. **(A)** Frequency sweep demonstrating the relatively consistent gel-like response regardless of Tm. **(B)** Flow sweep demonstrating the relatively consistent shear-thinning behavior regardless of Tm. **(C)** Elastic storage moduli (G') and viscous loss moduli (G'') at 1% strain and 10 rad/s of liposomal hydrogels with varying Tm. **(D)** Tan(delta) at 1% strain and 10 rad/s of liposomal hydrogels with varying Tm. All measurements were conducted at room temperature.

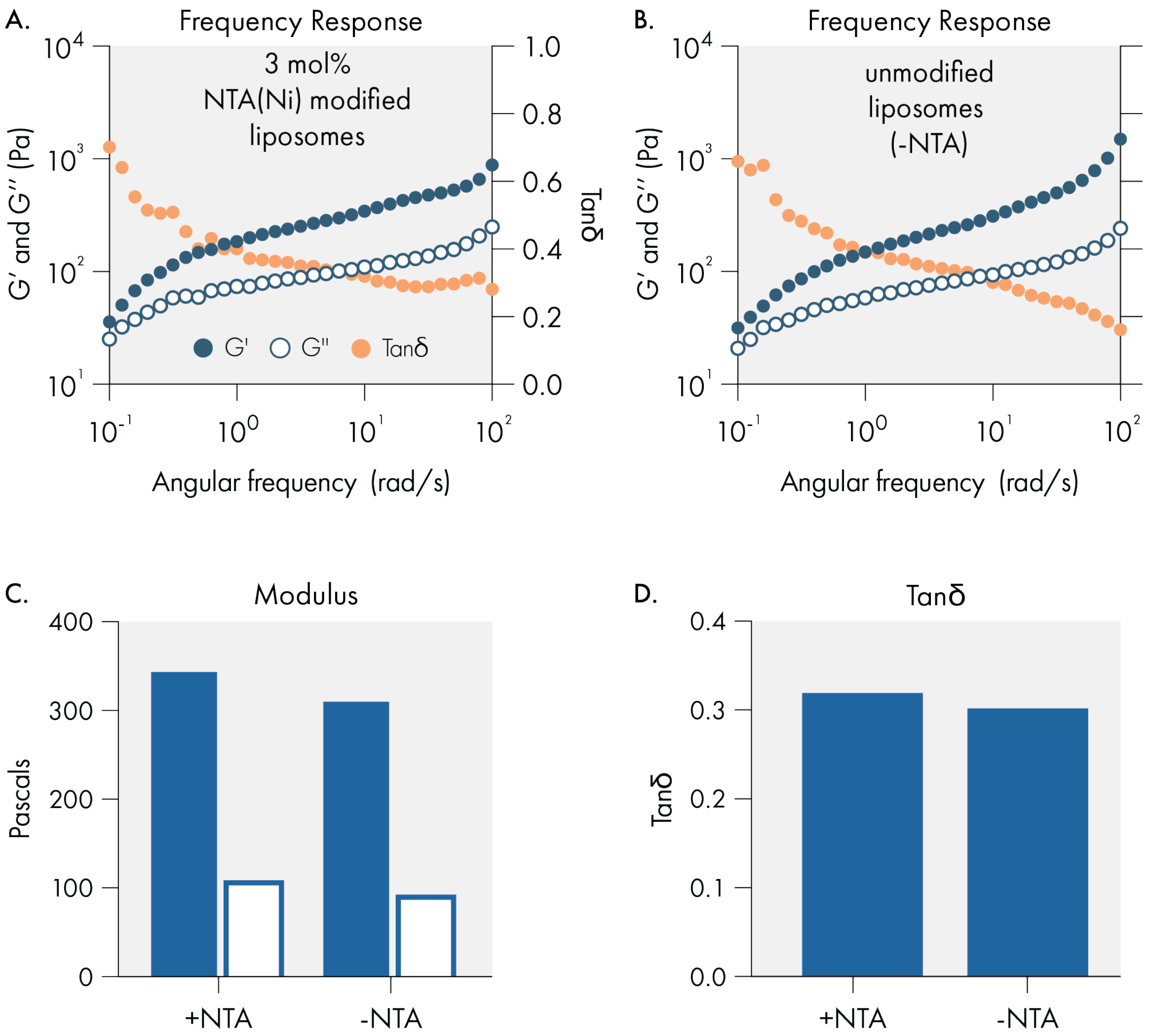

**Supplemental Figure 6**. Effect of 3 mol% NTA modified lipid incorporation on the rheological properties of liposomal hydrogels. No differences are observed. **(A)** Frequency sweep demonstrating the gel-like response. **(C)** Elastic storage moduli (G') and viscous loss moduli (G'') at 1% strain and 10 rad/s of both liposomal hydrogels. **(D)** Tan(delta) at 1% strain and 10 rad/s of both liposomal hydrogels. Both formulations were loaded with GFP to a final concentration 100 µg/mL. Rheological measurements were conducted using 8mm serrated parallel plates.

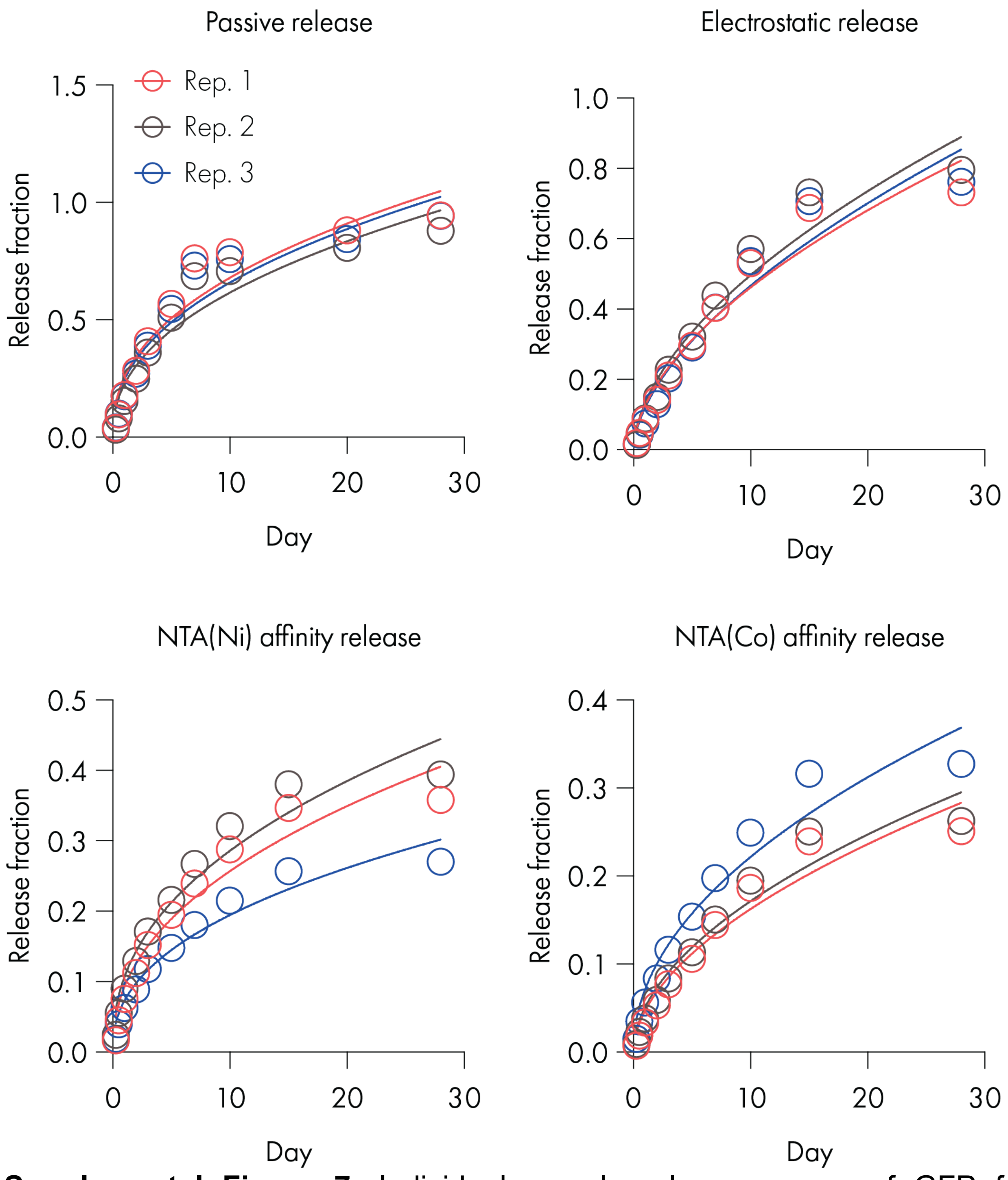

**Supplemental Figure 7.** Individual sample release curves of GFP from replicate hydrogels for several formulations of liposomal hydrogels that release cargo through different mechanisms with nonlinear Korsemeyer-Peppas fits.[1] Passive release is conducted with unmodified liposomes (Formulation 1 in Table 1) and GFP with no his tag. Electrostatic release is conducted with unmodified liposomes and GFP with a his tag. NTA(Ni) affinity release is conducted with hydrogels composed of 3 mol% NTA(Ni) modified liposomes (Formulation 8 in Table 1) and GFP with a his tag. NTA(Co) affinity release is conducted with hydrogels composed of 3 mol% NTA(Ni) modified liposomes (Formulation 9 in Table 1) and GFP with a his tag.

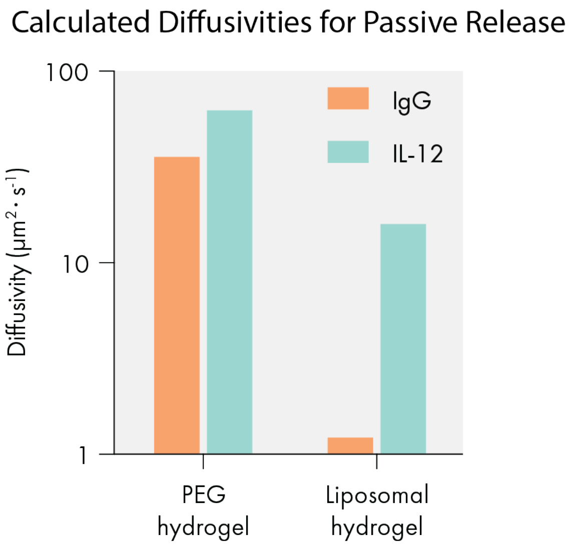

**Figure 8.** Predicted diffusivities of IgG antibodies and IL-12 cytokines in standard covalent PEG hydrogels and in liposomal hydrogels, assuming no matrix interaction with cargo (*e.g.*, passive release mechanisms).[2,3]
